# Cardiovascular risk in ANCA-associated vasculitis: monocyte phenotyping reveals distinctive signatures between serological subsets

**DOI:** 10.1101/2024.01.16.575967

**Authors:** Yosta Vegting, Katie ML Hanford, Aldo Jongejan, Gayle RS Gajadin, Miranda Versloot, Nelly D van der Bom-Baylon, Tamara Dekker, E Lars Penne, Joost W van der Heijden, Eline Houben, Frederike J Bemelman, Annette E Neele, Perry D Moerland, Liffert Vogt, Jeffrey Kroon, Marc L Hilhorst

## Abstract

**Objectives:** Anti-neutrophil cytoplasmic antibodies (ANCA)-associated vasculitides (AAV) is associated with an increased cardiovascular risk, particularly the myeloperoxidase AAV serotype (MPO-AAV). Distinct alterations in monocyte phenotypes may cause accelerated atherosclerotic disease in AAV.

**Methods:** A cohort including 43 AAV patients and 19 healthy controls were included for downstream analyses. Extensive phenotyping of monocytes and monocyte-derived macrophages was performed using bulk RNA-sequencing and flow cytometry. An *in vitro* transendothelial migration assay reflecting intrinsic adhesive and migratory capacities of monocytes was employed. Subsequent sub-analyses were performed to investigate differences between serological subtypes.

**Results:** Monocyte subset analysis showed increased classical monocytes during active disease, whereas non-classical monocytes were decreased. RNA-sequencing revealed upregulation of distinct inflammatory pathways and lipid metabolism-related markers in monocytes of active AAV patients. No differences were detected in the intrinsic monocyte adhesion and migration capacity. Monocytes of MPO-AAV patients in remission expressed genes related to inflammation, coagulation, platelet-binding and interferon signalling, whereas the expression of chemokine receptors indicative of acute inflammation and monocyte extravasation (i.e., CCR2 and CCR5) was increased in monocytes of proteinase-3(PR3)-AAV patients. During active disease, PR3-AAV was linked with elevated serum CRP and increased platelet counts compared to MPO-AAV.

**Conclusion:** These findings highlight changes in monocyte subset composition and activation, but not in the intrinsic migration capacity of AAV monocytes. MPO-AAV monocytes are associated with sustained upregulation of inflammatory genes, whereas PR3-AAV monocytes exhibit chemokine receptor upregulation. These molecular changes may play a role in elevating cardiovascular risk as well as in the underlying pathophysiology of AAV.

**Key messages:** - Monocytes are activated during active ANCA-associated vasculitis (AAV) and upregulate lipid metabolism-related markers

- AAV monocytes have a normal intrinsic adhesion and migration capacity, although overall monocyte migration likely rises by other mechanisms

- The two serological subsets MPO-AAV and PR3-AAV exhibit differences in monocyte activation and chemokine receptor expression

## Introduction

Risk of cardiovascular disease (CVD) is heightened with 65% in patients with anti-neutrophil cytoplasmic antibodies (ANCA)-associated vasculitides (AAV). CVD has emerged as the leading cause of death in these patients (1, 2). AAV encompass a group of autoimmune diseases characterized by systemic inflammation of small-to-medium sized blood vessels (3). ANCA are autoantibodies targeting proteinase 3 (PR3) or myeloperoxidase (MPO) found in granules of neutrophils and monocytes. These autoantibodies play a key role in the pathophysiology of AAV by activating the innate immune system resulting in local necro-inflammation and vital organ damage (4).

The origins of accelerated atherosclerotic disease in AAV are likely multifaceted, involving traditional CVD risk factors such as diabetes mellitus (5), and disease-specific risk factors such as endothelial damage, (glucocorticoid) therapy and vascular inflammation (6). Innate immune cells, most importantly monocytes and macrophages, play pivotal roles in the initiation and acceleration of CVD through processes such as transmigration into atherosclerotic plaques, differentiation into lipid-laden foam cells, and via their release of proinflammatory mediators (7). Anti-inflammatory therapies targeting macrophage cytokines interleukin-1β and interleukin-6 have shown significant potential against CVD (8, 9).

To date, little is known about the direct contribution of monocytes in driving the increased cardiovascular risk of AAV patients. Certain studies emphasize an elevated cardiovascular risk within the MPO-AAV subtype compared to its PR3-AAV counterpart (10, 11), while others did not (5, 12, 13). Even in remission, some studies indicate sustained monocyte activation and upregulation of adhesion markers in AAV (14, 15). Since CVD risk is linked to arterial wall inflammation (16), monocyte transendothelial migration and extravasation into plaques are likely to be key events in the pathogenesis of high cardiovascular risk of AAV patients as well. Targeting monocyte activation and their subsequent migration could be effective strategies for preventing cardiovascular disease in AAV.

The aim of this study was to investigate monocyte activation and the intrinsic migration capacity of monocytes of AAV patients, during both active disease and in remission.

## Methods

More detailed information on methods can be found in the Supplementary Data S1.

### Study subjects, samples and clinical information

Patients with AAV and controls were included. Blood samples were collected either once or twice, during active disease or in remission, defined by a Birmingham Vasculitis Activity Score version 3 (BVASv3) disease activity score of no more than one (17). Patients with a concurrent infection were excluded. On the day of blood withdrawal, information regarding demographic details, disease characteristics, and immunosuppressive medication use was collected and a standard laboratory panel was used to evaluate kidney function, a complete blood count and the presence of inflammation. Also, disease activity and treatment or vasculitis-related organ damage was measured by the BVASv3 and Vasculitis Damage Index (VDI) (18), respectively. The blood lipid profile, including total cholesterol, low-density lipoprotein (LDL), high-density lipoprotein (HDL), and triglycerides, and use of statin and antihypertensive therapy was retrospectively collected.

### PBMC isolation peripheral blood

Using standard density gradient centrifugation, peripheral blood mononuclear cells (PBMCs) were isolated from the blood and directly used for monocyte isolation or stored in liquid nitrogen.

### Bulk sequencing CD14^+^ bead selected monocytes and monocyte-derived macrophages

Bulk RNA sequencing was performed of CD14^+^-bead selected monocytes from patients with active (n=4) and stable disease (n=10) or healthy controls (n=6). Using freshly isolated PBMCs, CD14^+^ cells were positively selected by use of column-based immunomagnetic cell separation. Next, we performed bulk RNA sequencing of monocyte-derived macrophages (MDMs) from patients with active (n=1) and stable disease (n=3) and healthy controls (n=3). For macrophage culturing, 25 million PBMCs were thawed, plated for an hour and then washed to remove non-adherent cells. Adherent monocytes were cultured for five days with macrophage colony-stimulating factor (M-CSF, 50 ng/ml). On day five, MDMs were either stimulated with 10 ng/µl LPS and 50 ng/µl IFNy, 50 ng/µl IL4, 50 ng/µl IL-10, or left untreated for 24 hours. RNA was isolated and extracted after an on-column DNAse treatment. mRNA capture and library preparation was performed and cDNA was sequenced using paired-end sequencing, with a maximum read length of 150 bp on the Novaseq 6000 (Illumina) at a targeted sequencing depth of 40M reads/sample.

### RNA sequencing data processing and analysis

Excess adapter sequences, if present, were trimmed and reads were aligned to the human reference genome. Counts were obtained and quality control was performed. Genes with more than 2 reads in one or more samples were kept. Count data were transformed to log2-counts per million (logCPM), normalized, and precision weighted using voom (19). Differential expression was assessed using an empirical Bayes moderated t-test within limma’s linear model framework (20) including the precision weights. Adjusted *P*-values below 0.1, in consideration of the small sample size, were regarded statistically significant.

Geneset enrichment analysis (GSEA) was performed using CAMERA (limma package) with a value of 0.01 for the inter-gene correlation, using selected geneset collections (Hallmark collection and the BioCarta, KEGG and Reactome subsets of the C2 collection) from the Molecular Signatures Database (MSigDB; v2023.1.Hs). P-values were calculated using a two-sided directional test (direction of change, ‘up’ or ‘down’) and corrected for multiple testing using the Benjamini-Hochberg FDR. An adjusted *P*-value <0.1 was considered significant. Results for selected genesets and comparisons were visualized using enrichment networks.

Transcription factor (TF) activity was inferred using the decoupleR R package. The TF activity scores for the chosen comparison were inferred using the Univariate Linear Model method for TFs with at least 5 targets.

The main analysis was performed using R v4.1.0 (Bioconductor v3.13), the TF activity inference analysis was performed using R v4.3.0 (Bioconductor v3.17).

### Flowcytometric analysis of monocytes

Flowcytometric analysis of a panel related to monocyte adhesion and migration was performed. Cryopreserved PBMCs from ANCA patients with active disease (n=10), follow-up samples during stable disease (n=5) and age-and-sex matched healthy controls (n=5) were thawed. Samples from active AAV patients included equal subsets with and without concurrent high-dose (defined as ≥20 mg/day) glucocorticoid therapy. For each sample, six panels of markers related to monocyte adhesion and migration were measured (**Supp. Table 1**). For each staining reaction, in total 0.25 x 10^6^ PBMCs were incubated in a fixed staining volume (50ul) using pre-titrated antibodies. To mitigate batch effects, we incubated all samples with a pre-made antibody cocktail and conducted measurements on the same day. Gating strategy is shown in **Supp. Fig. 1**. To correct for background fluorescence, the Median Fluorescence Intensity (MFI) was subtracted with the MFI of the unstained backbone panel. A minimum cell count threshold of 200 cells was established as a criterion for inclusion in (MFI) analyses. A Wilcoxon rank-sum test was conducted to compare the expression of migration markers on peripheral blood monocyte subsets between ANCA patients and healthy controls. Principle component analysis (PCA) was used to summarize expression of markers and to highlight differences between monocyte subsets based on their marker profile. The marker values were log-transformed to stabilize variance and improve normality.

### Transendothelial monocyte migration assay

To assess the adhesive and migratory capacity of CD14^+^ monocytes, a transendothelial monocyte migration assay (TEM) was performed. Human Aortic Endothelial Cells (HAECs) were cultured in duplo or in triplo in a 12-well plate until reaching confluency and stimulated with 1 ng/mL IL-1β and 10 ng/mL TNFα overnight for 18 hours. HAECs were washed and media was changed at the day of the assay for at least 6 hours. Next, 100,000 viable CD14^+^-bead selected monocytes were loaded onto the HAECs and incubated for 30 minutes. Subsequently, adhered and migrated monocytes were fixed, imaged, and quantified. To control for bias, the outcome assessor was blinded for patient group.

## Results

### Patient characteristics

In total, 43 AAV patients with active and stable disease, and 19 HC healthy controls (HC) were included. Follow-up samples of 11 active AAV patients were collected after remission was achieved. Blood samples were used for blood marker analysis, monocyte and monocyte-derived macrophage phenotyping and a monocyte TEM assay. An overview of patient and disease characteristics for the total cohort can be found in Table 1 and **Suppl. Table S2-6** for an overview and characteristics of included patients per experiment. The majority of ANCA patients had renal involvement and were treated with immunosuppressive medication at time of blood withdrawal.

**Table 1:** Baseline characteristics patients total cohort. SD = standard deviation; IQR = interquartile range; BVAS = Birmingham Vasculitis Activity Score version 3; VDI = Vasculitis Damage Index; CRP = C-reactive protein; BMI = Body Mass Index; eGFR = estimated glomerular filtration rate; *Follow-up samples of 11 active patients were included in the remission group.

|  | Included ANCA patients (n=43)* |  | Healthy controls (n=19) |
| --- | --- | --- | --- |
|  | Active samples (n=24) | Remission samples (n=19)<br>Follow-up samples (n=11) |  |
| **Demographics** |  |  |  |
| Age, years, mean (SD) | 62 (12) | 61 (12) | 48 (21) |
| Female sex, n (%) | 9 (38) | 7 (23) | 12 (63) |
| BMI, mean (SD) | 27 (7) | 28 (9) | 24 (5) |
| **Disease characteristics** |  |  |  |
| **Clinical subtype** |  |  |  |
| \| GPA/MPA/eGPA/Renal limited, n | 18/3/1/2 | 24/4/0/2 | - |
| **ANCA subtype** |  |  |  |
| \| PR3/MPO, n | 19/5 | 22/8 | - |
| **Organ involvement** |  |  |  |
| \| Renal, n (%) | 17 (71) | 28 (93) | - |
| \| Pulmonary, n (%) | 14 (58) | 14 (47) | - |
| \| Cardiovascular, n (%) | 1 (4) | 3 (10) | - |
| \| ENT, n (%) | 10 (42) | 15 (50) | - |
| **Disease duration in days, median (IQR)** | 10 (1229) | 1181 (3846) | - |
| **eGFR, (CKD-EPI), per ml/min/1.73 m2, median (IQR)** | 41 (48) | 49 (41) | 96 (27) |
| **BVAS, median (IQR)** | 14 (7.25) | 0 (0) | - |
| **VDI, median (IQR)** | 0 (2) | 2 (2.75) | - |
| **CRP, mg/l, median (IQR)** | 30.1 (62.9) | 1.8 (3.1) | 1.5 (1) |
| **Monocytes, 10<sup>9</sup>/L, median (IQR)** | 0.60 (0.50) | 0.55 (0.29) | 0.50 (0.23) |
| **Use of immunosuppressive medication, n (%)** | 16 (67) | 22 (73) | 0 (0) |
| \| Glucocorticoid use, n (%) | 11 (49) | 14 (47) | 0 (0) |
| \| Prednisolone dosage users, mg, median (IQR) | 60 (1215) | 5 (0) | - |
| \| Methylprednisolone <30 days, n (%) | 6 (25) | 0 (0) | 0 (0) |
| \| Azathioprine, n (%) | 0 (0) | 4 (13) | 0 (0) |
| \| Cyclophosphamide, n (%) | 3 (13) | 0 (0) | 0 (0) |
| \| Rituximab < 1 year, n (%) | 5 (21) | 15 (50) | 0 (0) |
| **Cardiovascular risk** |  |  |  |
| **History of cardiovascular event, n (%)** | 2 (8) | 4 (13) | 0 (0) |
| **Diabetes mellitus, n (%)** | 3 (13) | 4 (13) | 2 (11) |
| **Current smoker, n (%)** | 3 (13) | 2 (7) | 1 (5) |
| **Total cholesterol, mmol/L, median (IQR) (n=23)** | 5.7 (1.7), n=7 | 5.1 (1.0), n=17 | NA |
| \| LDL, mmol/L, median (IQR) | 3.6 (2.0), n=8 | 3.0 (1.3), n=16 | NA |
| \| HDL, mmol/L, median (IQR) | 1.3 (0.4), n=7 | 1.4 (0.2), n=17 | NA |
| **Triglycerides, mmol/L, median (IQR)** | 1.6 (0.1), n=7 | 1.3 (0.6), n=17 | NA |
| **Use of statins, n (%)** | 7 (29) | 10 (33) | NA |
| **Use of antihypertensive medication, n (%)** | 11 (46) | 21 (70) | NA |

### Increased classical monocyte subset and decreased non-classical monocyte subset in AAV

To investigate possible origins of accelerated atherosclerotic disease in AAV, we first measured total peripheral blood monocyte and platelet count. AAV patients with active and stable disease had higher total monocyte and platelet count than controls (**Fig. 1a-b**). Next, we performed flow cytometric analysis and classified monocytes based on CD14 and CD16 expression into CD14^++^/CD16^−^ (Classical, CM), CD14^++^/CD16^+^ (intermediate, IM) and CD14^+^/CD16^++^ (Non-Classical, NCM) monocytes. The classification of monocyte subsets was validated through the assessment of established markers (21)(CM: high CCR2, IM: high CCR5, NCM: high CX3CR1, **Supp. Fig. 2**). Interestingly, compared to controls, the percentage of classical monocytes was significantly elevated during active disease (median % CM, 91.0 vs. 82.7, *P* = 0.0013), whereas the absolute number and percentage of non-classical monocytes was markedly decreased in active AAV patients (median % NCM, 5.8 vs. 13.0, *P* = 0.0087) (**Fig. 1c-d**). Likewise, bulk RNA sequencing of CD14^+^ bead-selected monocytes of MPO-AAV patients confirmed a significant reduction in the expression of NCM markers (**Fig. 1e**) (gene, log2FoldChange(log2FC); *FCGR3A/CD16,* −1.90*; LYPD2,* −7.06; *CX3CR1,* −2.32; *PECAM1,* −0.82; all markers adj. *P* < 0.01) compared to HC. Interestingly, the transcription factor *NR4A1*, important in the regulation of CM to NCM transition (22), was significantly decreased (log2FC −1.86, adj. *P* = 0.075). These findings may suggest a halt in monocyte transition, however, there was no substantial reduction observed in the inferred transcription factor *NR4A1* activity (**Fig. 1f**).

**Fig. 1:**
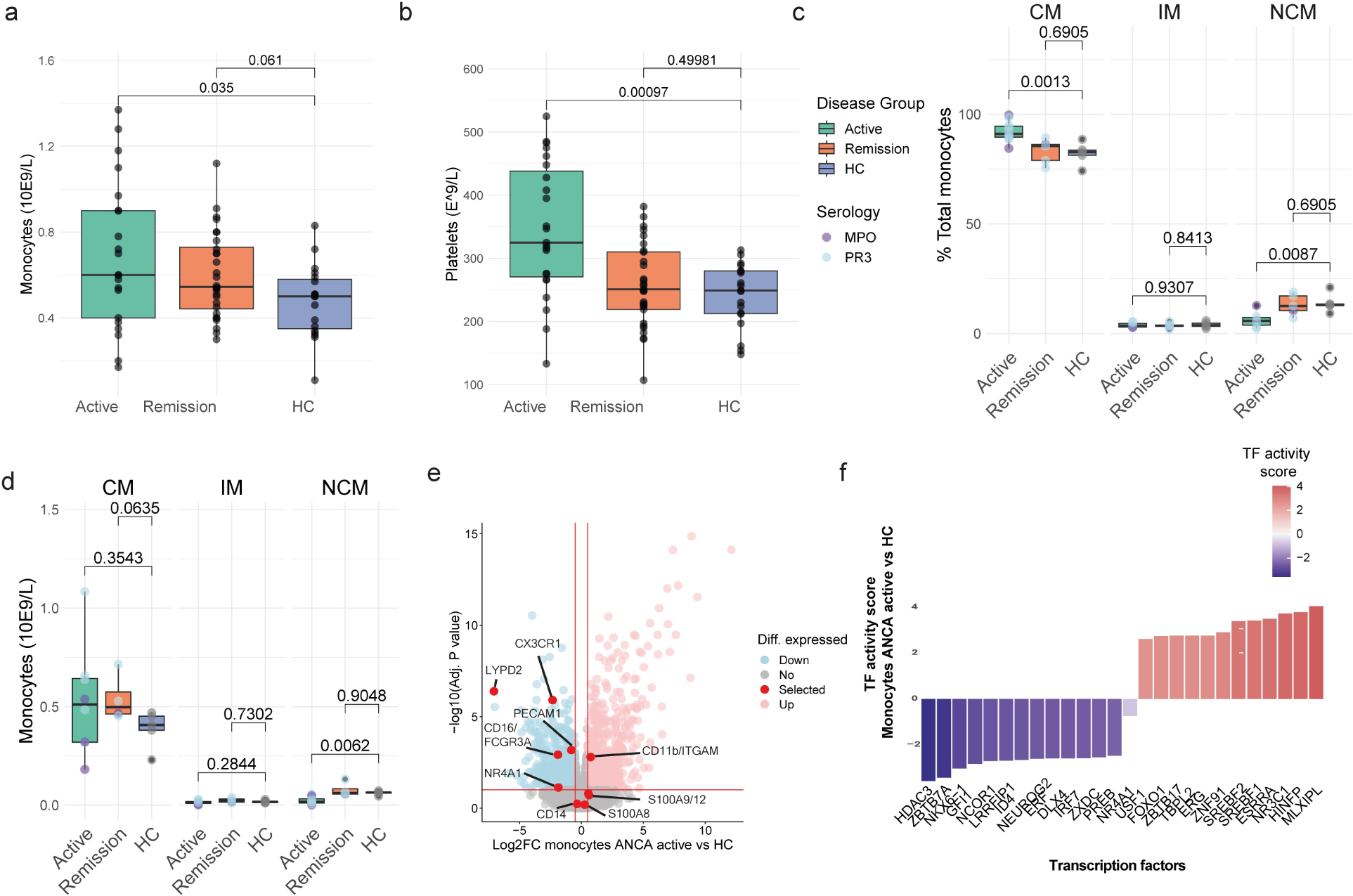
Changes in monocyte subset composition in AAV. **a,** Boxplot showing peripheral blood monocyte count total cohort. **b,** Boxplot showing blood platelet count total cohort. **c,** Boxplot showing peripheral blood monocyte subset count in FACS cohort. **d**, Boxplot showing percentage of peripheral blood monocyte subsets. **a-d,** Statistics were calculated using a Wilcoxon rank-sum test. **e,** Volcano plot of differentially expressed genes related to monocyte subsets. Monocytes of ANCA patients with active disease (n=4) and healthy controls (n=6) were isolated and gene expression was analysed using bulk mRNA sequencing. Cut-offs of a log2FoldChange of >=0.5 and adj. *P* <0.1 are marked with a red line. **f**, Bar plot showing transcription factor (TF) activity score per transcription factor monocytes active AAV vs HC. CM, Classical monocyte; IM, Intermediate monocyte; NCM, non-classical monocyte.

### Proinflammatory monocyte activation with altered lipid metabolism profile

RNA sequencing data was further analyzed by a CAMERA-based geneset enrichment analysis (GSEA) using four MSigDB geneset collections (Hallmark, BioCarta, KEGG, Reactome). In remission, no enrichment in inflammatory signaling pathways was found if monocytes of MPO-AAV and PR3-AAV patients were pooled (**Fig. 2a**). However, monocytes of active MPO-AAV patients more highly expressed multiple gene sets associated with IL1 signaling, peroxisome and NFκB activation, while interferon α response was decreased (**Fig. 2b**). Additionally, gene sets associated with lipid processing and metabolism, such as cholesterol biosynthesis (**Supp. Data S1**), and lipid markers, like *PLIN2*, *LPL*, *PPARγ* were significantly upregulated (**Fig. 2c**) indicating an altered lipid metabolism profile. Also, mRNA expression of complement receptor *C3AR1* and anti-inflammatory macrophage markers such as *CD163*, *C1QC*, and *MRC1* were significantly upregulated. Interestingly, mRNA expression of the complement receptor C5AR1, recently identified as a novel treatment target in ANCA (23), did not show significant alterations (logFC 0.59, adj. *P* = 0.23). A selection of these markers was validated by flow cytometry but did not confirm differences on the protein levels (**Fig. 2d, Supp. Fig. 3**). Since all active AAV monocyte samples received glucocorticoid treatment, a subanalysis was performed (**Fig. 2e, Supp. Fig. 6**) and showed significant elevation of C3AR1 expression of CM of AAV patients with high-dose glucocorticoid treatment (*P* = 0.03), but not in patients without. Since serum and urine CD163 are biomarkers of disease activity (4), and samples with highest *CD163* expression were from treatment-naïve patients, monocytic *CD163* upregulation is likely disease-specific.

**Fig. 2:**
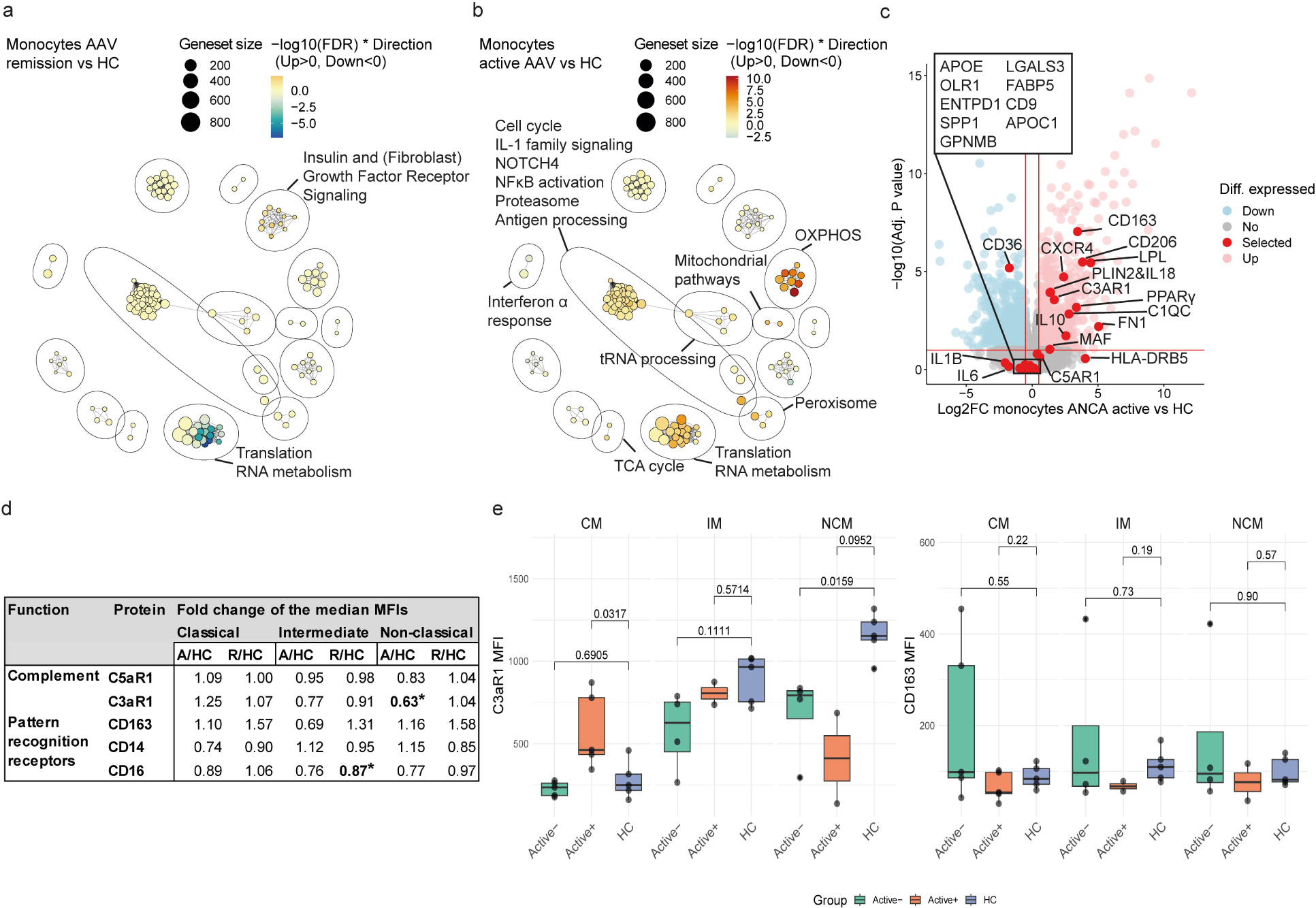
Proinflammatory monocyte activation with altered lipid metabolism profile during active disease. **a-b,** Geneset enrichment analyses (GSEA) of monocytes using four MSigDB geneset collections (Hallmark, BioCarta, KEGG, Reactome). Enrichment networks are shown to group significant (FDR<0.1) genesets in all comparisons based on genes they have in common. The nodes represent genesets that are significantly upregulated (red) or downregulated (blue). Overlap between genesets is represented by lines. Annotation was manually added based on significance. **a,** GSEA monocytes AAV patients in remission (n=10) compared to HC (n=6). **b,** GSEA monocytes active AAV patients (n=4) compared to HC (n=6). **c,** Volcano plot of differentially expressed genes related to markers of monocyte activation and lipid metabolism. Cut-offs of a log2FoldChange of >=0.5 and adj. *P* <0.1 are marked with a red line. **d,** Fold changes of the median MFIs of surface markers determined by FACS between active (A) AAV patients (n=10), follow-up samples in remission (R) (n=5) and matched HC (n=5). * *P* < 0.05. **e,** Boxplots showing differences in MFI of C3aR1 and CD163 between patients treated with (Active+) and without (Active-) high dosages of glucocorticoids. **d-e,** Statistics were calculated using a Wilcoxon rank-sum test. A, active; R, remission; HC, Healthy control; AAV, ANCA-associated vasculitis; CM, Classical monocyte; IM, Intermediate monocyte; NCM, non-classical monocyte.

### Monocyte-derived macrophages (MDMs) of PR3-AAV patients do not retain their inflammatory profile in remission

Prolonged inflammatory stimuli can induce long-term changes in the responsiveness of innate immune cells, a process which is called trained immunity (24). To test whether AAV monocytes retain their inflammatory profile after differentiation into macrophages, we performed bulk RNA sequencing of *in vitro* cultured MDMs with and without 24h of proinflammatory (LPS and IFNy) or anti-inflammatory (IL4 or IL10) stimulation. Interestingly, the success rate of macrophage cultures with AAV samples was limited compared to controls (samples with RNA yield > 1000ng per stimulation; AAV vs. HC, 48% vs. 69%). This discrepancy is likely attributed to increased cell mortality as a consequence of prolonged activation of these monocytes. Therefore, at time of sequencing, only one active PR3-AAV sample was available, limiting our analyses. First, inflammasome-related cytokine expression was examined in these cells (**Fig. 3a**). Whereas *IL-18* and *IL-10* expression were significantly (Log2FC *Il-18* 1.39, adj. *P <*0.001; *IL-10* 2.57, adj. *P* = 0.019) elevated in monocytes of active MPO-AAV patients(**Fig. 2c**), no significant differences were found in unstimulated and LPS+IFNy stimulated MDMs of the active PR3-AAV patient (**Fig 3b**) and of the PR3-AAV patients in remission (n=3)(**Fig. 3c**). Also, GSEA and transcription factor activity inference did not show upregulation of gene sets and key transcription factors related to inflammation (25) (**Fig. 3d, Supp. Data S1**), excluding an inflammatory cell fate in PR3-AAV.

**Fig. 3:**
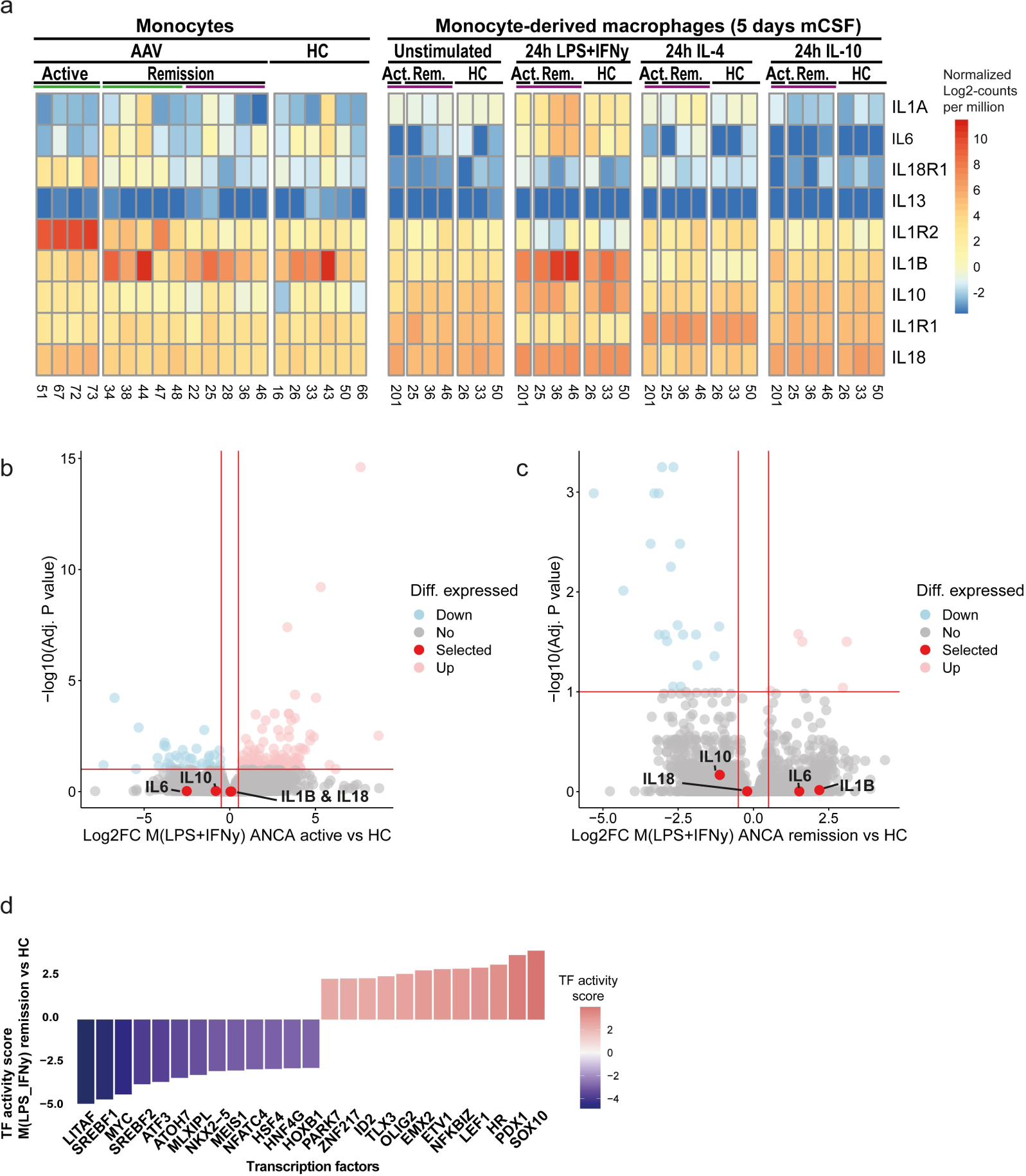
Monocyte-derived macrophages of PR3-AAV patients do not retain their inflammatory profile in remission. **a,** Heatmap of expression of selected interleukins and receptors of monocytes and monocyte-derived macrophages(MDMs). Green lines represent MPO-AAV patients, purple PR3-AAV patients. Each column represents an individual patient sample. **b-c,** Volcano plot of differentially expressed genes related to proinflammatory cytokine production in MDMs stimulated with 24h LPS+INFγ. Cut-offs of a log2FoldChange of >=0.5 and adj. *P* <0.1 are marked with a red line. **b,** M(LPS+INFγ) active AAV patient (n=1) vs HC (n=3). **c,** M(LPS+INFγ) AAV patients in remission (n=3) vs HC (n=3). **d**, Bar plot showing transcription factor (TF) activity score per transcription factor M(LPS+INFγ) AAV patients in remission vs HC. HC, healthy control.

### No differences in the expression of leukocyte migration markers or intrinsic transendothelial migration capacity

Monocytes play a significant role in development of atherosclerosis by transmigration into atherosclerotic plaques (7). GSEA did not show increased expression of gene sets related to leukocyte migration during active and stable disease (**Supp. Data S1**). Individual marker analyses showed increases in mRNA expression of *ITGAM (*CD11b) and *CXCR4*, while *ITGAL* (CD11a)*, ITGA4 (*CD49D)*, PECAM1, CX3CR1* and *CCR5* were significantly decreased in monocytes of active AAV patients compared to healthy controls (**Fig. 4a**). Flow cytometric analysis did not confirm corresponding up- or downregulation of these markers (**Fig. 4b, Supp. Fig S3-5**). No clear difference in the expression of other chemokine receptors and adhesion molecules between those with and without high-dose glucocorticoids was found (**Supp. Fig S7-8**). To assess the intrinsic adhesive and migratory capacity, an *in vitro* TEM assay was performed. No differences in intrinsic migration capacity were found between ANCA patients and healthy controls (**Fig. 4c**). However, within active patients, distinct differences in monocyte adhesion and migration were observed between patients treated with high-dose corticosteroids and untreated individuals, suggesting a strong inhibitory effect of prednisone on monocyte migration.

**Fig. 4:**
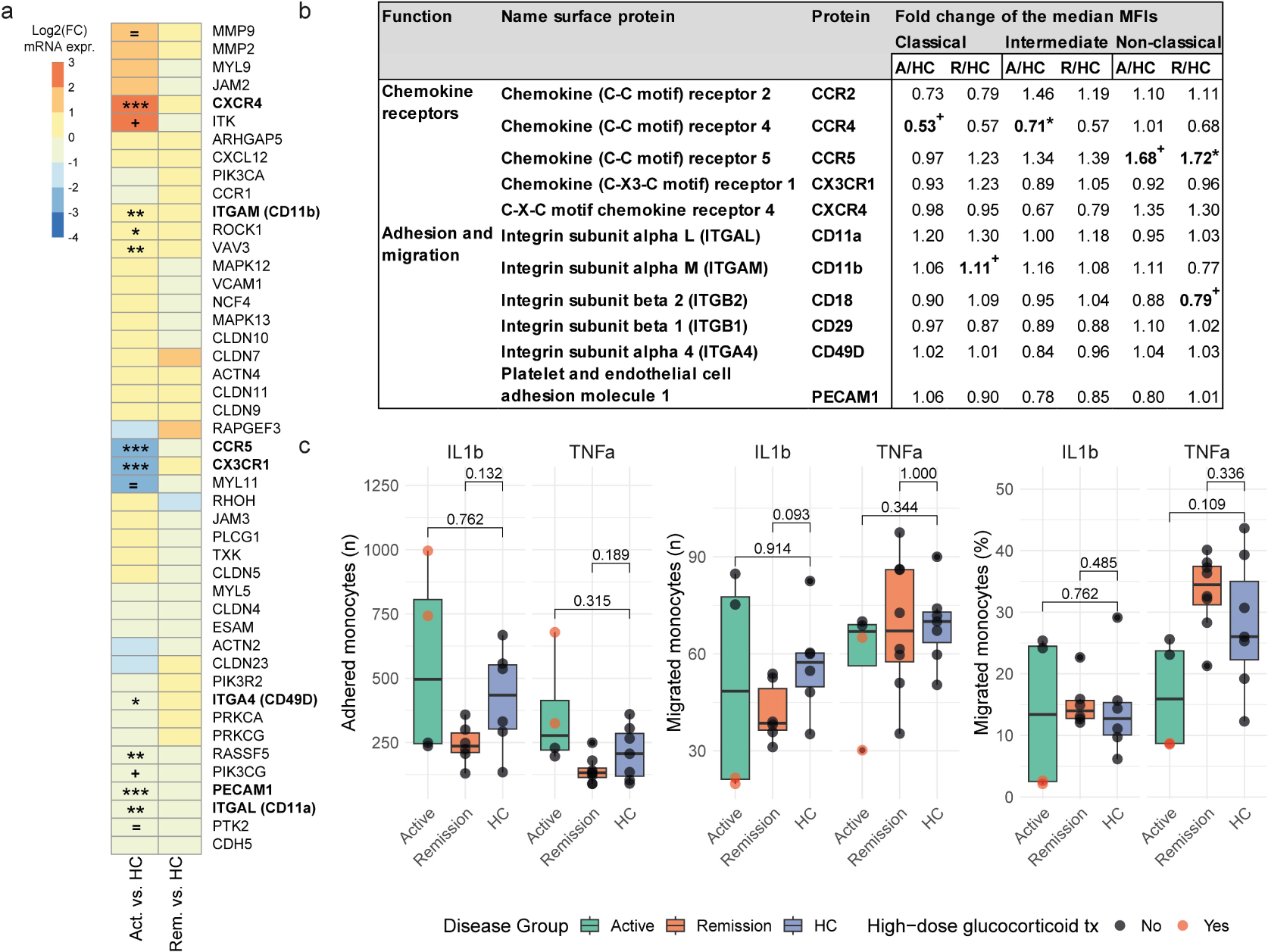
No differences in the intrinsic monocyte adhesion and migration capacity. **a,** Differences in mRNA expression of molecules related to migration between monocytes of ANCA patients with active disease (n=4), remission (n=10), and healthy controls (n=6). Individual markers from a gene set related to transendothelial migration (KEGG) and based on the literature (52) are shown. Genes that were further studied by flow cytometric analysis are highlighted in bold. Adjusted P-values are indicated. **b,** Fold changes of the median MFIs of surface markers determined by FACS between active AAV patients (n=10), follow-up samples in remission (n=5) and matched HC (n=5). Wilcoxon rank-sum test was performed to compare the expression between groups within monocyte subsets. **c,** Boxplot showing monocyte adhesion and migration of isolated CD14 positive monocytes to TNF-α or IL1β overnight-stimulated Human Aortic Endothelial Cells (HAECs). Wilcoxon rank-sum test was performed to compare between groups. High-dose prednisone treatment is shown in purple. C, Classical monocyte; IM, Intermediate monocyte; NCM, non-classical monocyte. P values, = *P* = 0.1 - 0.25, + *P* = 0.05 - 0.1, \**P* < 0.05, ** *P* < 0.01, *** *P* < 0.001.

### MPO-AAV monocytes in remission express genesets related to inflammation and monocyte-platelet activation, while PR3-AAV monocytes upregulate chemokine receptors CCR2 and CCR5

To investigate differences between monocytes obtained from MPO and PR3-AAV patients, we performed sub-analyses of the earlier described experiments. No statistical differences in total peripheral blood monocyte (subset) counts within active or remission were found, although highest classical monocyte counts were found in PR3-AAV patients (**Fig. 5a**). Since only samples of active MPO-AAV patients were included for monocyte bulk RNA sequencing, no comparisons were made for the active disease group. Within the remission patients, MPO-AAV had increased expression of gene sets related to inflammation (**Fig. 5b**). Comparing between the two serotypes, PR3-AAV monocytes had lower expression of gene sets related to inflammation, platelet and complement activation, coagulation and an interferon signature (**Fig. 5c**). Similarly, using transcription factor (TF) activity inference, monocytes of PR3-AAV patients during remission showed decreased activation of TFs involved in monocyte and macrophage differentiation (26, 27) (C/EBPβ), interferon and inflammatory signalling (NF-κB, FOS) (25, 28, 29), Il4/13 mediated macrophage activation (STAT6) (30). Also, TFs involved in glucocorticoid signalling (NR3C1) were downregulated in PR3-AAV monocytes, likely attributed to concurrent corticosteroid therapy in MPO-AAV patients (**Fig. 5d**). As shown by the GSAE, there was a decrease in expression of platelet activation markers (*TUBB1, ITGA2B, ITGB3, PF4, PPBP, SELP*) and markers of monocyte-platelet interaction (*SELPLG*) in monocytes of PR3-AAV compared to MPO-AAV patients during remission, although only *PF4* remained significant (Log2FC −2.48, adj. *P* = 0.073) after adjustment for multiple testing (**Fig. 5e, Supp. Data S1**). These data indicate the presence of monocyte-platelet aggregates in MPO-AAV, which are linked to CVD (31). Also, even with concurrent glucocorticoid treatment, monocytes from MPO-AAV patients exhibit a heightened proinflammatory state compared to their PR3 counterparts. Looking into migration markers, PR3-AAV monocytes during remission were associated with higher mRNA expression of *CCR2 (*Log2FC 0.80, adj. *P* = 0.071) and *CCR5 (*Log2FC 0.86, adj. *P* = 0.60) (**Fig. 5f**) compared to MPO-AAV, which was validated by flow cytometric analysis (**Fig. 5g-h, Supp. Fig. S9-11**), indicating increased responsiveness to chemotactic signals in PR3-AAV during remission. Due to limited MPO-AAV samples, statistical differences could not be tested in remission. *In vitro* monocyte adhesion and migration was not significantly different between PR3 and MPO-AAV patients in remission (**Fig. 5i**), although a large spread was found especially in PR3-AAV. Blood markers of inflammation showed higher platelet count, CRP and estimated glomerular filtration rate (eGFR) in PR3-AAV patients compared to MPO-AAV during active disease, while in remission no significant changes were found (**Fig. 5j**). To conclude, in remission, MPO-AAV monocytes express genesets related to inflammation, interferon signalling, coagulation, and monocyte-platelet activation, while adhesion markers and chemokine receptors are downregulated in comparison with PR3-AAV. Despite these changes in monocyte activation and chemokine receptor expression between serological subsets, monocyte migration remained unaffected in remission. During active disease, PR3-AAV is linked to increased markers of systemic inflammation, suggesting a different role for monocytes in these AAV subsets.

**Fig. 5:**
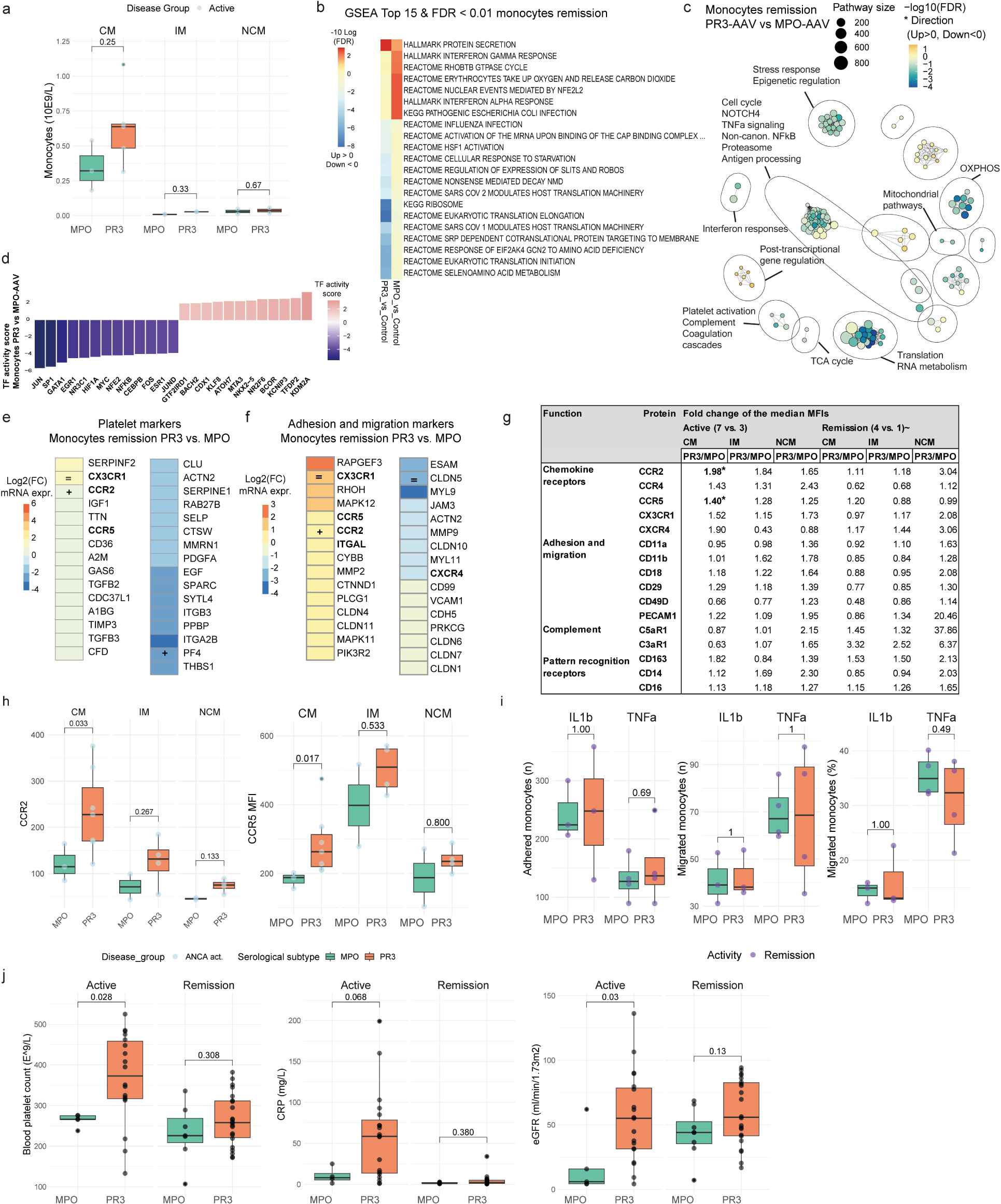
Differences in monocyte activation and migration between serological subtypes. **a,** Boxplot showing peripheral blood monocyte subset count in FACS cohort per serological subtype. Statistics were calculated using a Wilcoxon rank-sum test. **b-c,** Geneset enrichment analysis (GSEA) of monocytes using selected MSigDB geneset collections. Heatmap of top 15 significantly (FDR < 0.01) upregulated (red) and downregulated (blue) gene sets in monocytes PR3-AAV and MPO-AAV patients versus controls **c,** Enrichment networks are shown to group significant (FDR<0.1) genesets from selected comparisons based on genes they have in common. The nodes represent genesets that are significantly upregulated (red) or downregulated (blue) in PR3-AAV (n=5) compared to MPO-AAV (n=5) in remission. Overlap between genesets is represented by lines. Annotation was manually added based on significance. **d**, Bar plot showing transcription factor (TF) activity score per transcription factor PR3-AAV vs MPO-AAV monocytes in remission. **e-f,** Differences in mRNA expression of molecules related to migration (**e**) and platelet activation **(f)** between PR3-AAV vs MPO-AAV monocytes. Adjusted P values are indicated. Genes that were studied by flowcytometric analysis are highlighted in bold. **g,** Fold changes of the median MFIs of surface markers determined by FACS between active PR3-AAV (n=7) and MPO-AAV (n=3) patients, and between PR3-AAV (n=4) and MPO-AAV (n=1) patients in remission. **h,** Boxplots showing the MFIs of chemokine receptors; CCR2, CCR5. **g-h,** Wilcoxon rank-sum test was performed to compare the expression between groups within monocyte subsets. ∼ test could not be performed due to limited samples. **i,** Boxplot showing 30-min monocyte adhesion and migration of isolated CD14 positive monocytes to TNF-α or IL1β overnight-stimulated Human Aortic Endothelial Cells (HAEC) of MPO-AAV (n=4) and PR3-AAV (n=4) patients. Wilcoxon rank-sum test was performed to compare between groups. **j**, Boxplots showing peripheral blood platelet concentration, CRP and eGFR in total cohort per serological subtype. P values, = *P* = 0.1 - 0.25, + *P* = 0.05 - 0.1, \**P* < 0.05, ** *P* < 0.01, *** *P* < 0.001.

## Discussion

Accelerated atherosclerotic disease poses a significant health concern for patients with ANCA-associated vasculitis (AAV). The causes are likely multifactorial, but our understanding of the direct contribution of monocytes is currently limited. In this study, we found an increased activation profile of monocytes in AAV. No differences in the intrinsic migration capacity were found. Our data suggest that (sustained) monocyte activation in AAV might contribute to an accelerated cardiovascular risk.

Notably, we found decreased NCM differentiation, while the CM subset was increased in AAV compared to controls. There are mixed findings on monocyte composition in AAV (4), similar to the field of CVD (32). Whereas in most CVD studies an increased percentage of IM and NCM was found (32), in AAV, upregulation of CM appears intricately linked to the underlying pathophysiology of the disease, as shown by a recent single-cell RNA sequencing study, highlighting a link between the expansion of activated and interferon signature expressing classical monocyte subsets and resistance to therapy and kidney infiltration (33, 34). As expected, the CM subset expressed high CCR2 levels, and it was found that CCR2 was associated with arterial wall inflammation (35). Therefore, the increased CM subset is likely to play a significant role in both disease and cardiovascular risk, as evidenced by the heightened cardiovascular risk, particularly in the initial months before and after diagnosis (36, 37).

Next, bulk RNA sequencing of CD14^+^ bead-selected monocytes of AAV patients showed upregulation of genesets related to monocyte activation during active disease, and in MPO-AAV patients in remission. As CM constitute the primary monocyte subtype, these findings likely represent the activation of the classical monocyte subset. In CVD, monocytes exhibited increased release of proinflammatory cytokines (IL1β, IL6) after LPS or IFNγ treatment (38), similar to *in vitro* ANCA stimulation of healthy monocytes (4). In our study, unstimulated monocytes of active AAV patients expressed high IL-10 and IL18, whereas IL1β, IL6 levels were unchanged. Determination of an inflammatory cell fate lacks robustness, possibly caused by unintended selection of PR3-AAV samples with limited inflammation by loss of samples with persistent inflammation.

Further looking into characteristic features of atherosclerosis, the total panel of migration markers and intrinsic monocyte migration capacity were not enhanced in AAV patients compared to controls. Monocyte adherence and migration are increased in CVD, with upregulation of integrins (e.g. CD11b) (39, 40). Also, monocytes of active and stable AAV patients expressed increased integrin CD11b (14, 41). Prior to our study, no investigations into leucocyte extravasation had been conducted in AAV, leaving the functional implications of individual marker expression unknown until this point. Since the intrinsic migration capacity is unaffected, monocytes of AAV patients may potentially have similar migrating capacity into atherosclerotic plaques. However, in tissues with local inflammation, the release of monocyte-attracting chemokines (e.g. MCP-1) and inflammatory mediators is likely increasing net monocyte migration (42, 43). This is further supported by the observation that large numbers of (monocyte-derived) macrophages infiltrate into peripheral tissues including the kidneys (4).Therefore, increased monocyte attraction could also lead to increased net monocyte migration into atherosclerotic plaques, but more research remains necessary.

Platelet binding and activation is another driver for increased cardiovascular risk by recruitment of monocytes and the formation of platelet-monocyt complexes (44). We observed increased platelet numbers during active disease, which could enhance thrombus formation in combination with monocyte-platelet activation. Indeed, in AAV patients, an elevated platelet count was linked to the incidence of strokes (45). Also, the incidence of venous thromboembolic events (VTE) was significantly elevated and related to disease activity (46). Therefore, monocyte-platelet activation might be another risk factor for CVD in AAV.

A selection of studies, but not all, indicated a higher cardiovascular risk in the MPO-AAV subtype (10, 11). Strikingly, in remission, monocytes of MPO-patients expressed a more proinflammatory phenotype than its PR3-AAV counterparts. Also, monocytes of these patients upregulated markers related to platelet activation and monocyte-platelet aggregates, possibly contributing to a higher CVD risk during periods without apparent disease activity (31). MPO also directly modulates cardiovascular risk, since it is released locally by activated blood neutrophils in the formation of neutrophil extracellular traps (47) and serum MPO is an independent risk factor for CVD (48). MPO-AAV was associated with an increased risk of VTE (49), which may be caused by platelet activation. Interestingly, during active disease, different trends were seen. PR3-AAV was associated with a more inflammatory phenotype, with higher, but non-significant, (classical) monocyte levels, increased CCR2 and CCR5 expression on CM, in addition to elevated markers of inflammation in the blood, such as platelet count. CCR2 expression on circulating monocytes is correlated to monocyte extravasation (40) and inflammation of the arterial wall (35), and therefore directly linked to cardiovascular risk (16). Due to the limited sample size, we were unable to confirm *in vitro* increased monocyte migration in PR3-AAV during active disease and did not show differences in remission. However, immunostainings of AAV kidney biopsies show increased infiltration of classical MDMs in PR3-AAV patients compared to MPO-AAV (data submitted), which is likely caused by increased CCR2 expression on CM. These findings along with the clinical course suggest that PR3-AAV is associated with a more acute cardiovascular risk due to CCR2^+^ CM migration and activation specifically related to PR3-AAV disease flares, while monocyte-mediated inflammation in MPO-AAV might be more smouldering and related to platelet activation. As a result, MPO-AAV patients could be diagnosed in a later stage, reflected by a lower eGFR and signs of chronicity in these patients (50). Similarly, it could be speculated that low-grade inflammation in remission caused by MPO-AAV monocytes contributes to a long-term increased cardiovascular risk.

Our study highlights the intricate interplay between immune dysregulation and cardiovascular risk and may be important for (future) clinical patient care. Targeted monocyte therapies may have the potential to not only reduce cardiovascular risk but also directly mitigate the disease itself, offering dual benefits. Even with glucocorticoid treatment, MPO-AAV patients experienced ongoing monocyte activation in remission with signs of platelet binding and activation. Therefore, in these patients, anti-inflammatory therapy may potentially be intensified by addition of specific drugs, such as IL1β monoclonal antibodies. In addition, it might be interesting to assess whether initiating antiplatelet therapy at the time of diagnosis would provide more benefits than potential bleeding risks. Nevertheless, it remains important to focus on effective management of traditional risk factors (1, 6, 51).

Strengths encompass the use of freshly isolated samples in monocyte sequencing and migration experiments contributing to robustness of the data. The flow cytometry experiment was performed without batch variation and involved a thorough monocyte gating strategy, which was also validated by the expression of established markers (21). Limitations involve limited sample sizes due to logistical reasons and sample variability, including concurrent glucocorticoid therapy, posing challenges in interpreting findings as results may be influenced by therapy or the selection of patients with severe disease. Furthermore, the uneven distribution of ANCA serotypes across experiments made it difficult to investigate the differences between these subtypes effectively.

To conclude, we demonstrate an increased classical monocyte subset and increased monocyte activation in AAV compared to controls, without differences in the monocyte intrinsic migration capacity. MPO-AAV is associated with sustained low grade monocyte inflammation in combination with monocyte-platelet activation. On the other hand, PR3-AAV is related to more acute inflammation and increased classical monocyte chemotaxis due to CCR2 upregulation. These mechanisms could each individually contribute to an increased cardiovascular risk in AAV and enhance our understanding of the role of monocytes in the development of CVD.

## Data availability statement

Processed data are publicly available at ArrayExpress under accession number E-MTAB-13676 (https://www.ebi.ac.uk/biostudies/arrayexpress/studies/E-MTAB-13676?key=ceddcbb2-5cd5-47df-8bc0-40b1f895141f). The raw RNA human sequencing data that support the findings of this study are not publicly available due to privacy reasons and were therefore deposited into controlled access data storage from the European Genome-phenome Archive. These data are available from the corresponding author upon reasonable request (Data Access Committee: EGAC50000000024).

## Acknowledgements

We would like to thank Berend Hooibrink from Microscopy and Cytometry Core Facility AmsterdamUMC.

## Ethics statement

The research protocol was approved by the local ethics boards in accordance with the Declaration of Helsinki, and subjects gave their written informed consent.

## Supplementary information

**Supp. Fig. 1:**
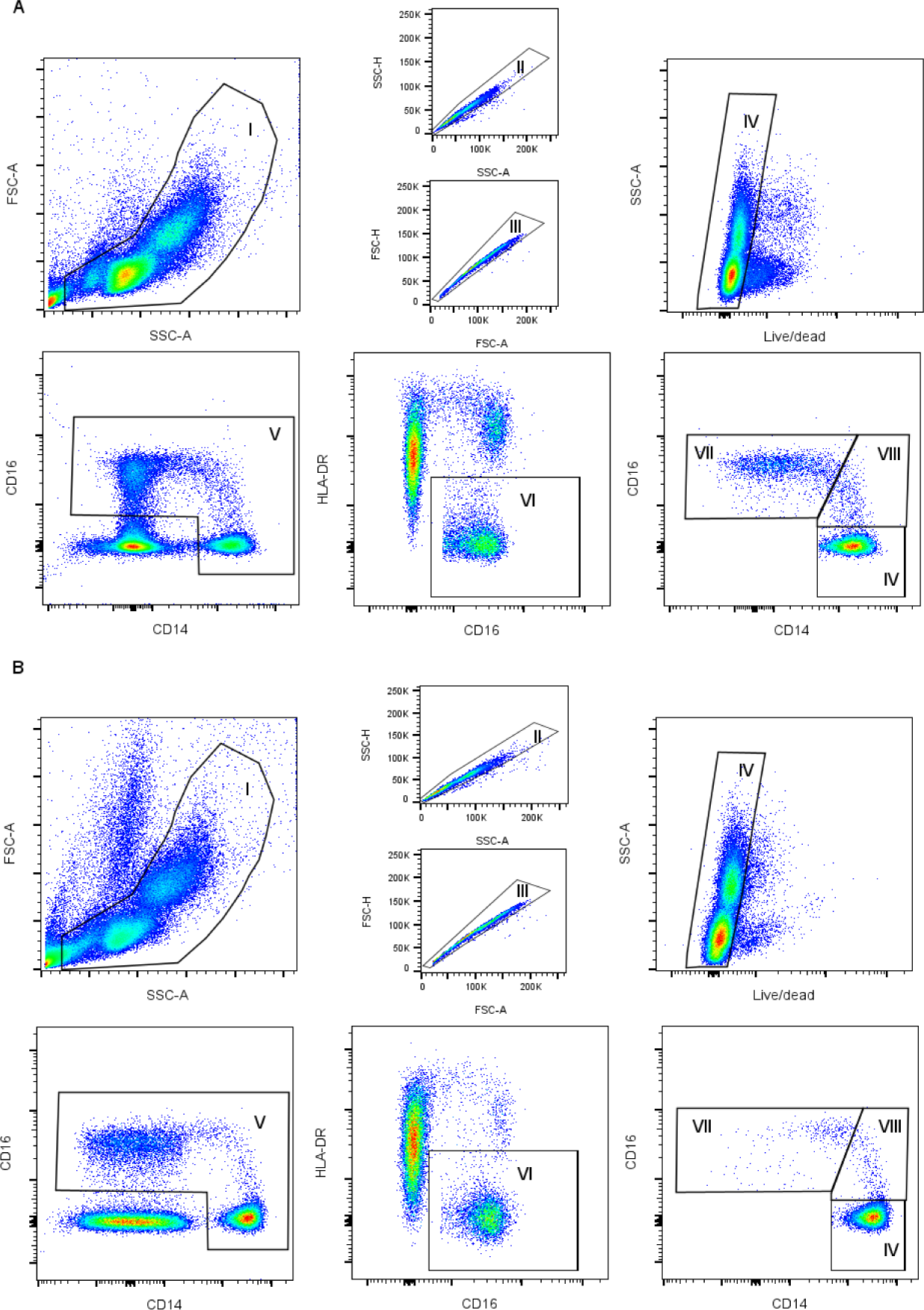
gating strategy. Gating of monocyte subtypes using an example of a healthy control (**A**) or AAV patient with active disease (**B**). First, Residual RBC, granulocytes and debris were excluded on the FSC/SSC plot (gate I). Next, a singlet gating was performed using a SSC-H/SSC-A (II) and FSC-H/FSC-A (gate III) plots. Viable cells were selected on the SSC-A/Live/dead plot (gate IV) and presented on a CD14/CD16 plot to select CD14^+^ and/or CD16^+^ cells (gate V). HLA-DR-cells were excluded (gate VI), followed by gating of monocyte subsets, CD14^+^/CD16^++^ (non-classical) (gate VI), CD14^++^/CD16^+^ (intermediate) (gate VIII), and CD14^++^/CD16^−^ (classical) (gate IV) monocytes.

**Supp. Fig 2.**
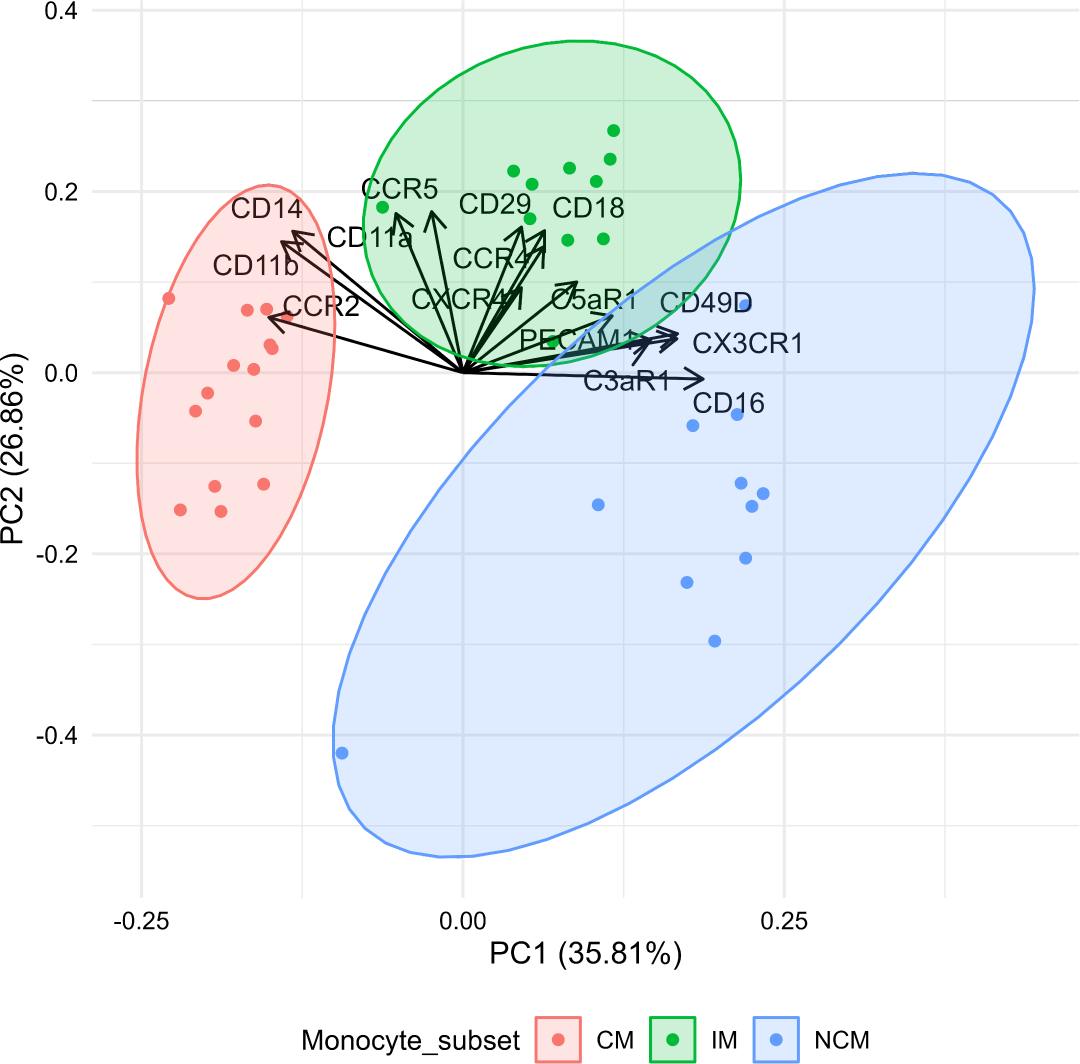
Validation of monocyte subset gating by specific monocyte surface marker expression. Principal Component Analysis (PCA) plot demonstrates the variance in surface marker profiles across monocyte subsets. Vectors indicate individual markers, with their direction and length indicating the trend and contribution to variance, respectively. Ellipses around each monocyte subset visualizes the variability within each monocyte subset. CM, classical monocytes; IM, intermediate monocytes; NCM, non-classical monocytes

**Supp. Fig 3.**
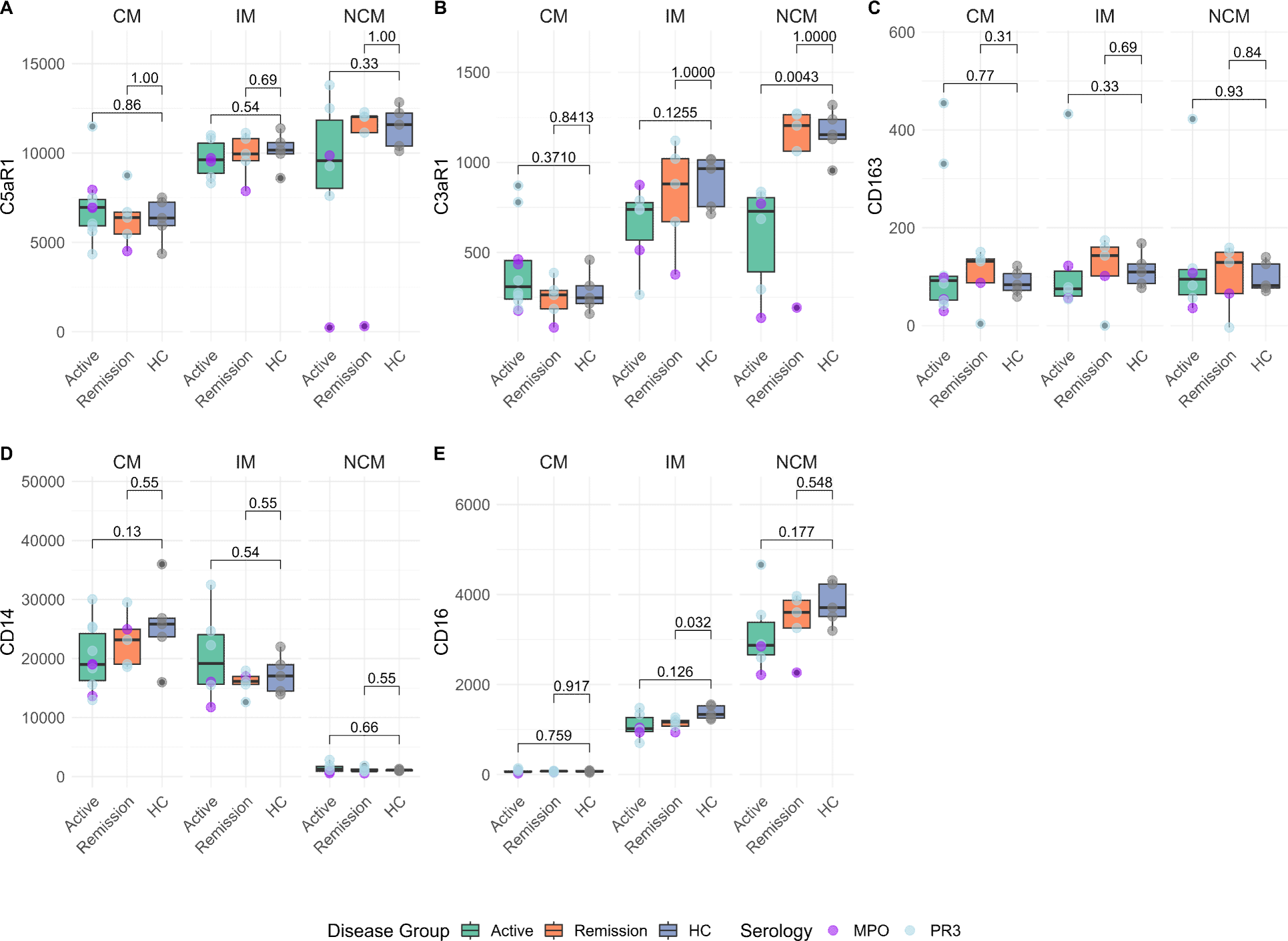
Expression of activation and differentiation markers. Boxplots showing the expression of activation and differentiation markers; A. C5AR1 B. C3AR1 C. CD163 D. CD14 E. CD16. Wilcoxon rank-sum test was performed to compare the expression between groups within monocyte subsets. Serological subtypes are depicted as MPO-ANCA (purple) and PR3-ANCA (blue).

**Supp. Fig 4.**
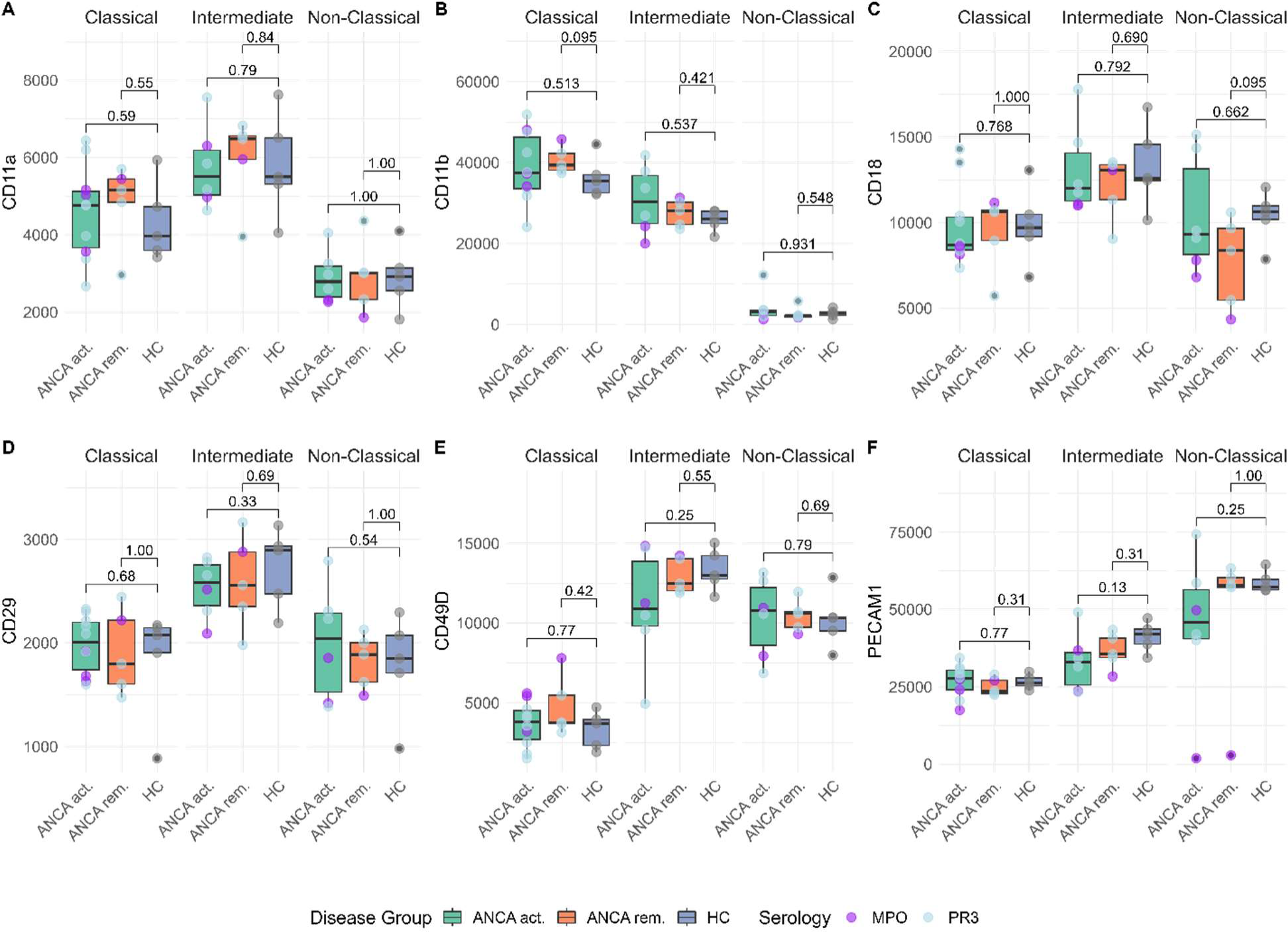
No clear differences in adhesion molecules and integrins between ANCA patients and healthy controls. Boxplots showing the expression of adhesion molecules; A. CD11a B. CD11b, C. CD18, D. CD29, E. CD49D, F. PECAM1. Wilcoxon rank-sum test was performed to compare the expression between groups within monocyte subsets. Serological subtypes are depicted as MPO-ANCA (purple) and PR3-ANCA (blue). Act. = active, rem = remission, HC= healthy controls

**Supp. Fig 5.**
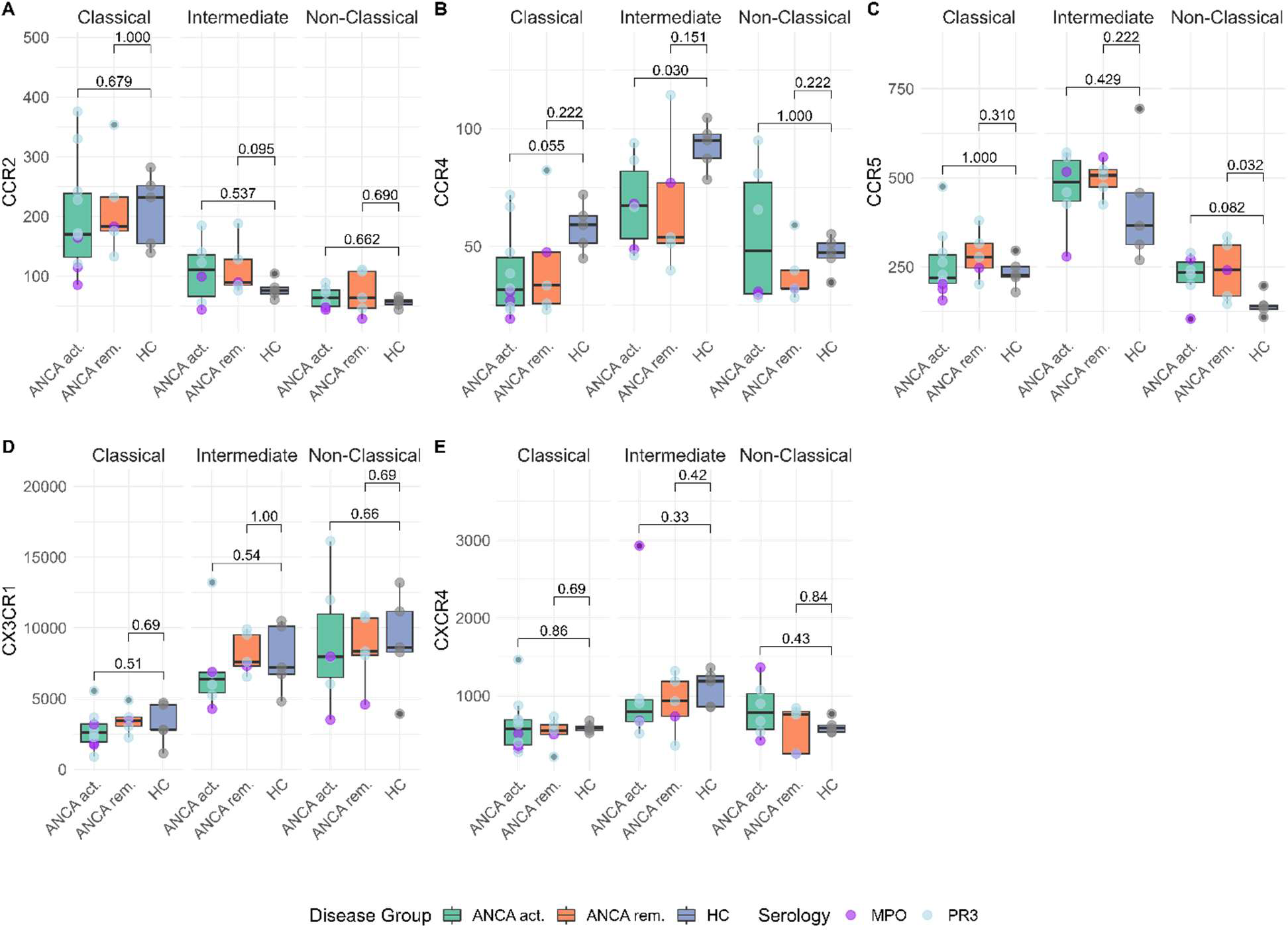
No clear differences in chemokine receptors between ANCA patients and healthy controls. Boxplots showing the expression of chemokine receptors; A. CCR2 B. CCR4, C. CCR5, D. CX3CR1, E. CXCR4. Wilcoxon rank-sum test was performed to compare the expression between groups within monocyte subsets. Serological subtypes are depicted as MPO-ANCA (purple) and PR3-ANCA (blue). Act. = active, rem = remission, HC= healthy controls

**Supp. Fig 6.**
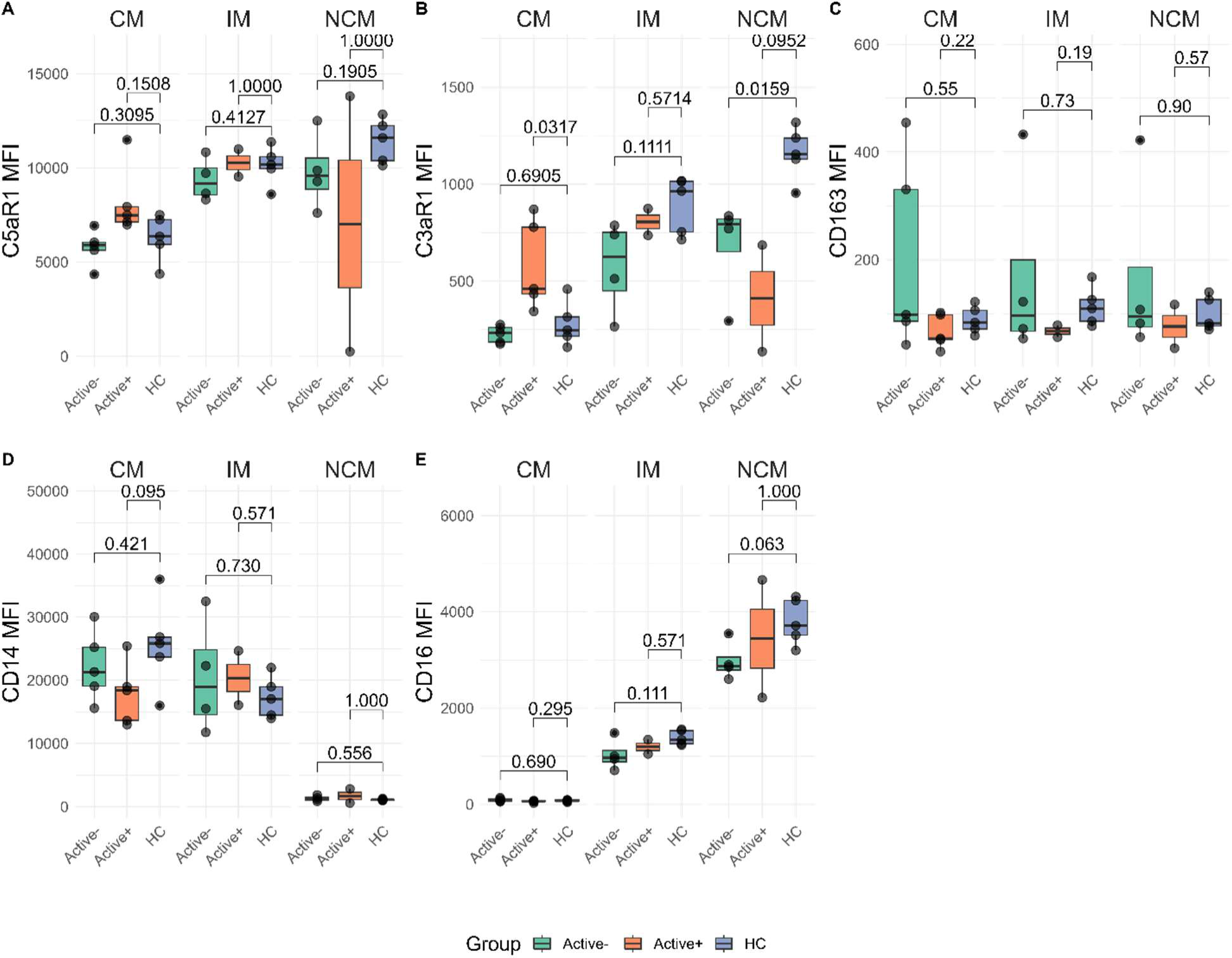
Expression of activation and differentiation markers divided on high dosage concurrent glucocorticoid treatment. Boxplots showing the expression of activation and differentiation markers; A. C5AR1 B. C3AR1 C. CD163 D. CD14 E. CD16. Wilcoxon rank-sum test was performed to compare the expression between groups within monocyte subsets. Active-, Active AAV patients without high-dose glucocorticoid treatment; Active+, Active AAV patients with high-dose glucocorticoid treatment; HC, healthy controls.

**Supp. Fig 7.**
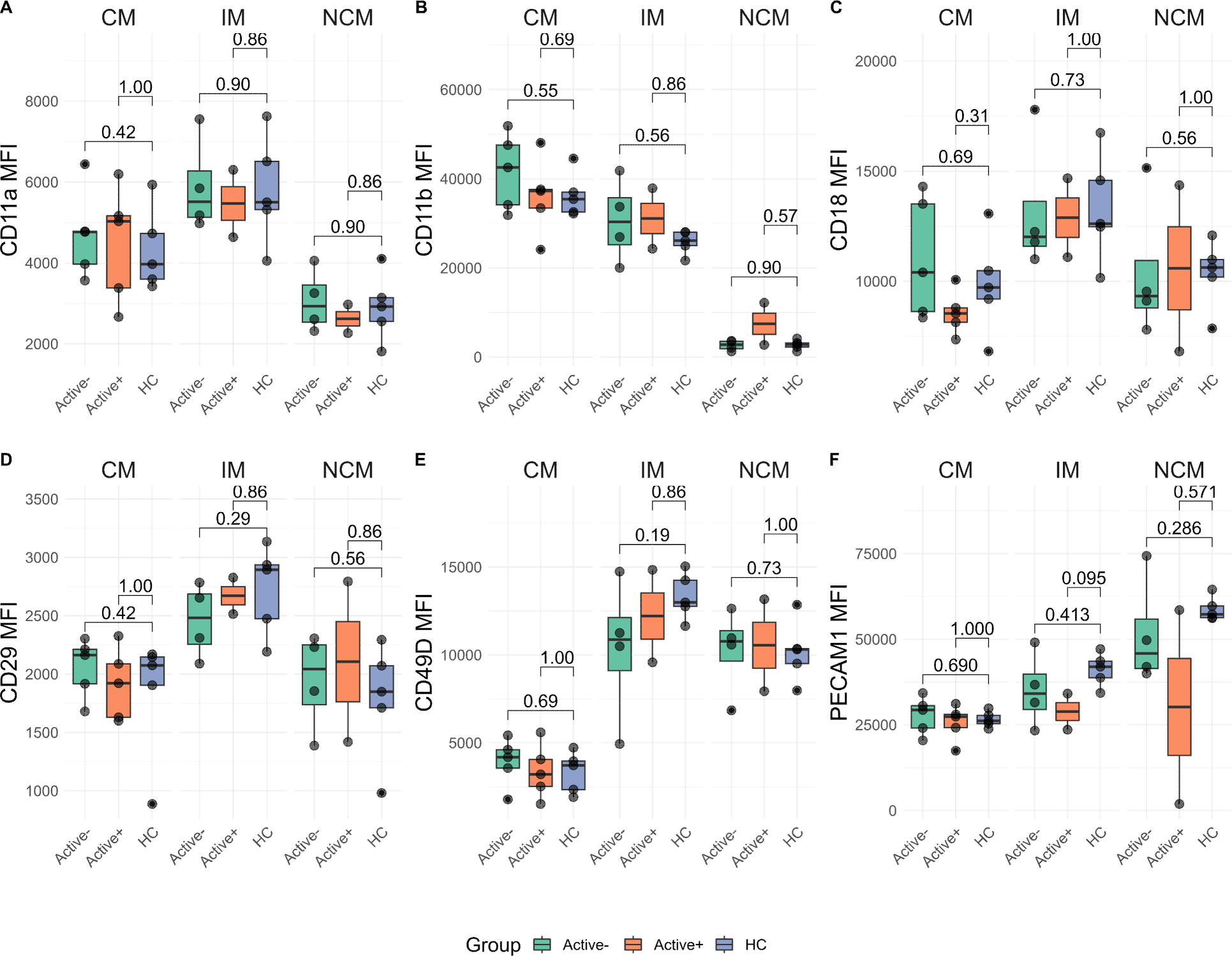
Expression of adhesion molecules and integrins divided on high dosage concurrent glucocorticoid treatment. Boxplots showing the expression of adhesion molecules; A. CD11a B. CD11b, C. CD18, D. CD29, E. CD49D, F. PECAM1. Wilcoxon rank-sum test was performed to compare the expression between groups within monocyte subsets. Active-, Active AAV patients without high-dose glucocorticoid treatment; Active+, Active AAV patients with high-dose glucocorticoid treatment; HC, healthy controls.

**Supp. Fig 8.**
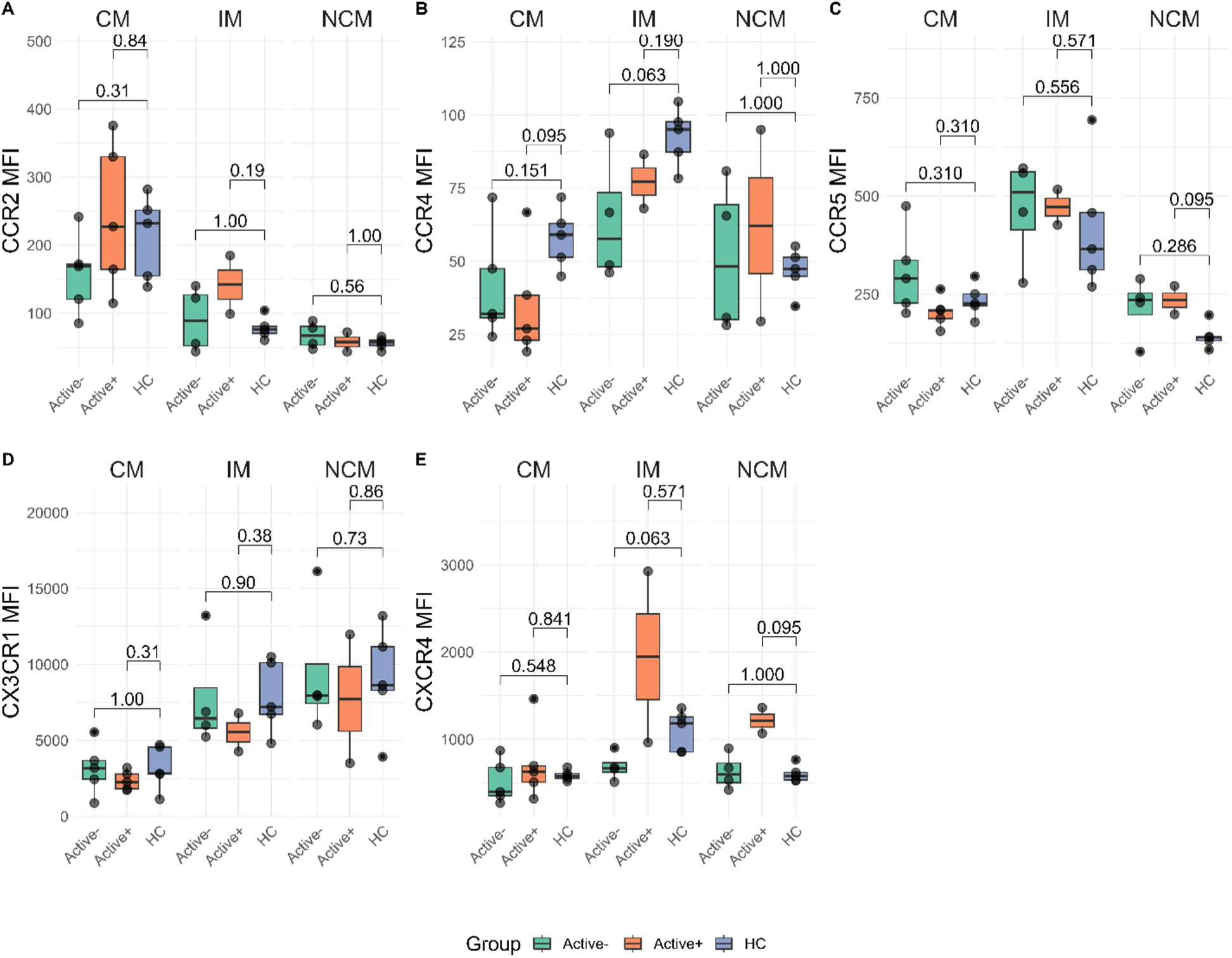
Expression of chemokine receptors divided on high dosage concurrent glucocorticoid treatment. Boxplots showing the expression of chemokine receptors; A. CCR2 B. CCR4, C. CCR5, D. CX3CR1, E. CXCR4. Wilcoxon rank-sum test was performed to compare the expression between groups within monocyte subsets. Active-, Active AAV patients without high-dose glucocorticoid treatment; Active+, Active AAV patients with high-dose glucocorticoid treatment; HC, healthy controls.

**Supp. Fig 9.**
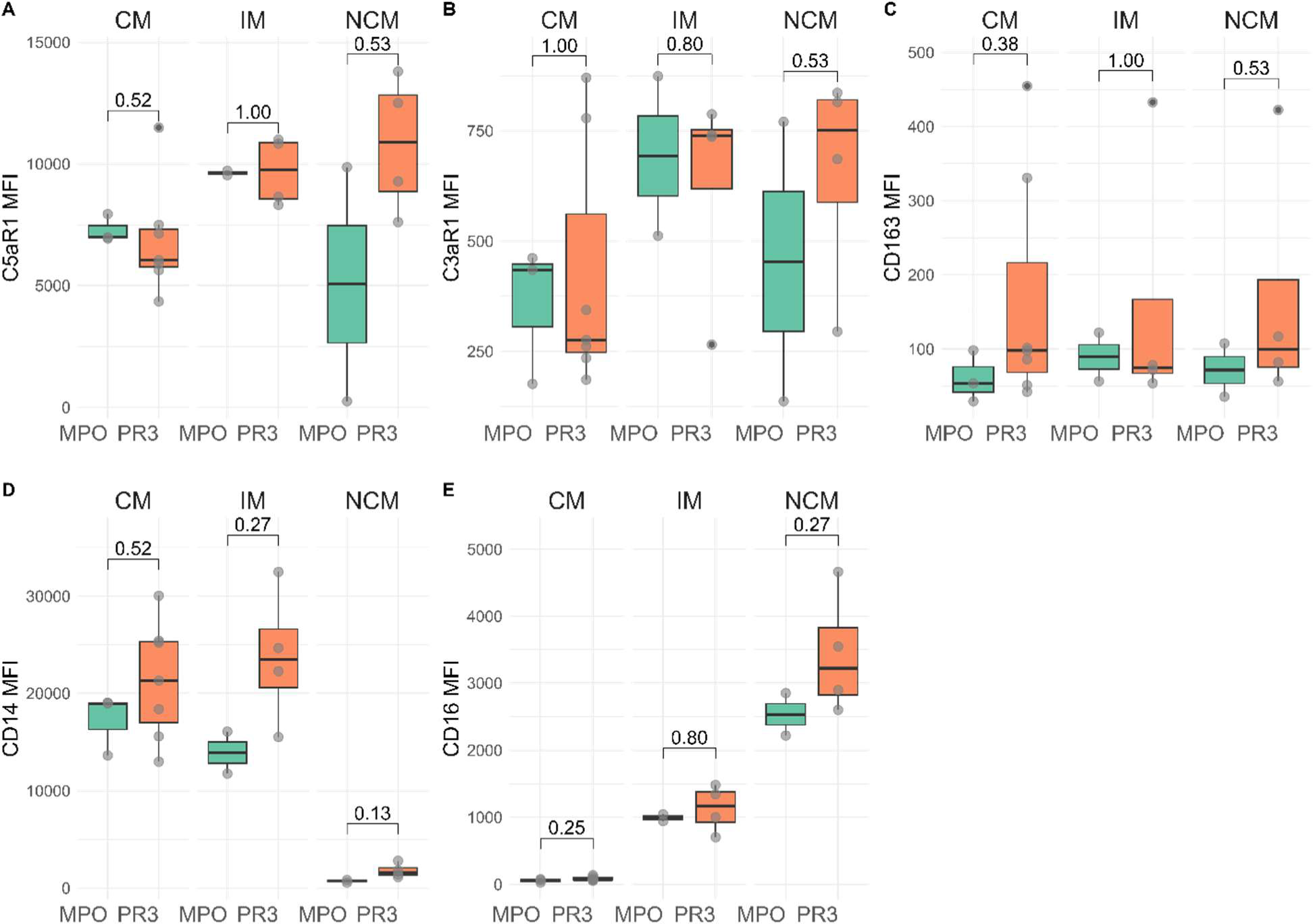
No differences in activation and differentiation markers between MPO-AAV and PR3-AAV patients. Boxplots showing the expression of C5aR1, C3aR1, CD163, CD14 and CD16. Wilcoxon rank-sum test was performed to compare the expression between serological subtypes within monocyte subsets. CM, classical monocyte; IM, intermediate monocyte; NCM, non-classical monocyte.

**Supp. Fig. 10:**
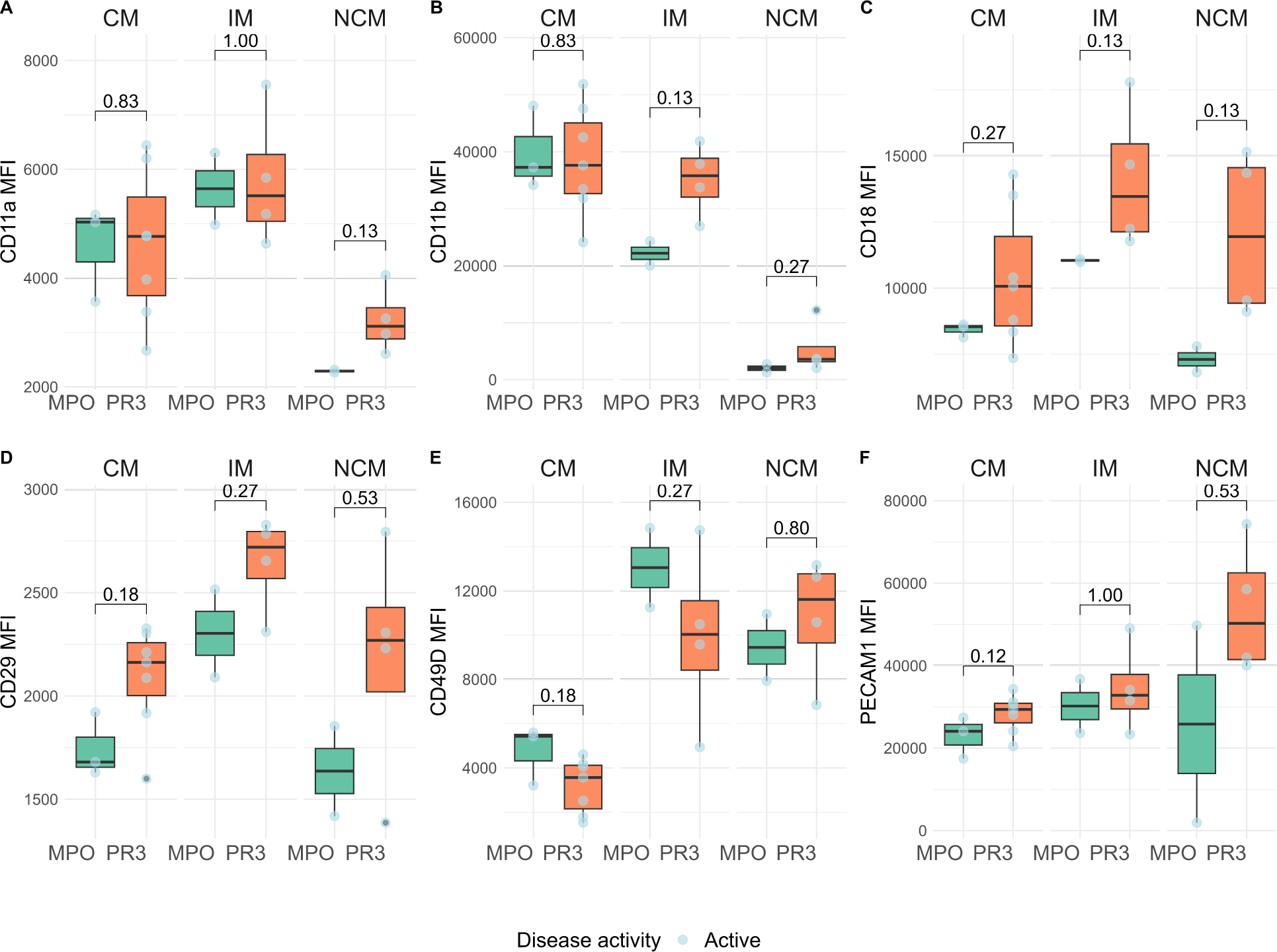
Differences in adhesion molecules and integrins during active disease between ANCA serological subtypes. Boxplots showing the expression of adhesion molecules; A. CD11a B. CD11b, C. CD18, D. CD29, E. CD49D, F. PECAM1. Wilcoxon rank-sum test was performed to compare the expression between groups within monocyte subsets. CM, classical monocyte; IM, intermediate monocyte; NCM, non-classical monocyte.

**Supp. Fig. 11.**
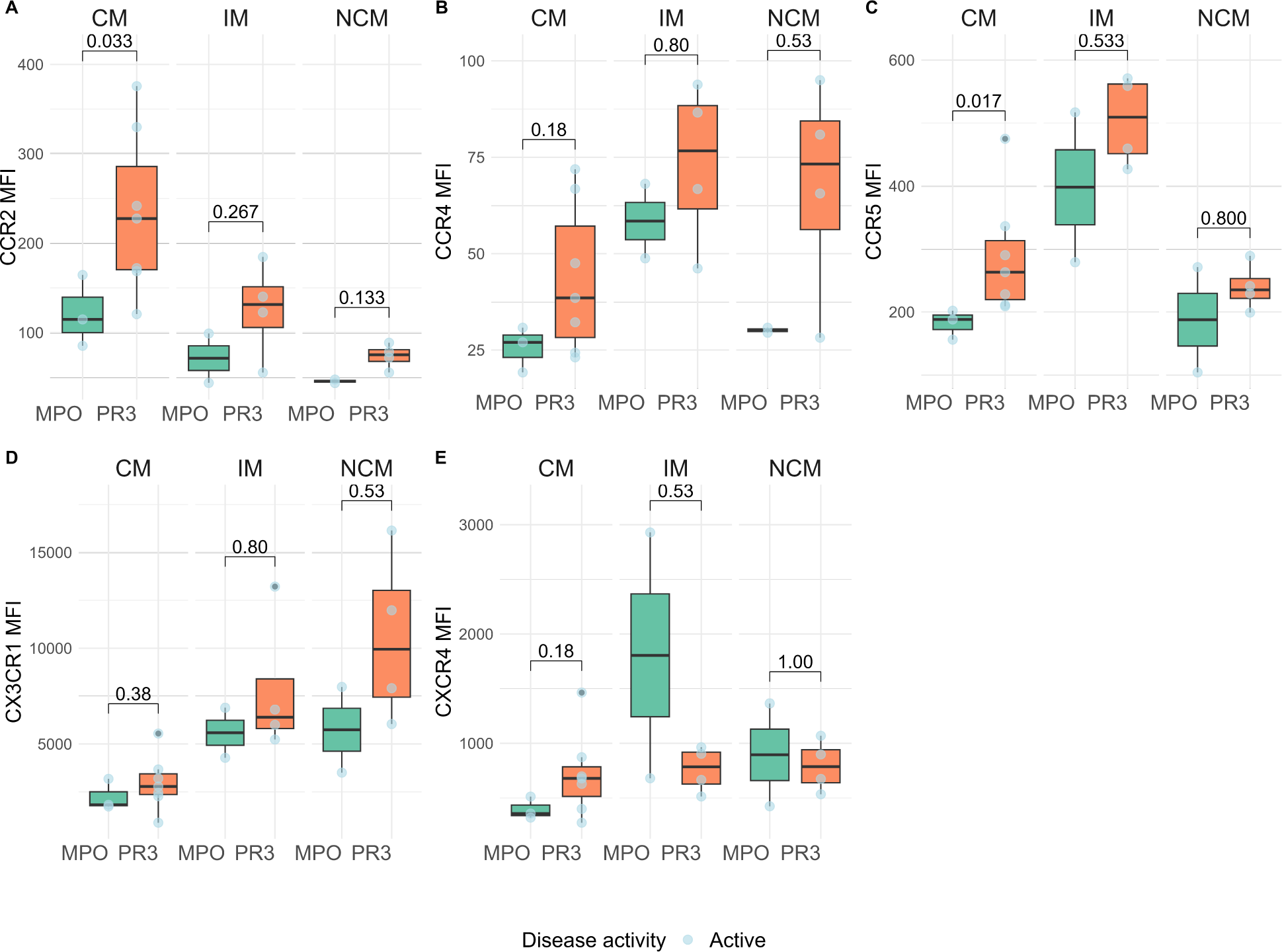
Upregulation of CCR2 and CCR5 on classical monocytes of PR3-AAV patients. Boxplots showing the expression of chemokine receptors; A. CCR2 B. CCR4, C. CCR5, D. CX3CR1, E. CXCR4. Wilcoxon rank-sum test was performed to compare the expression between groups within monocyte subsets. CM, classical monocyte; IM, intermediate monocyte; NCM, non-classical monocyte.

## Supplemental tables

**Supp. Table S1:** Monoclonal antibodies for flowcytometry.

| Antibody | Clone | Fluorochrome | Manufacturer | Product number | Dilution |
| --- | --- | --- | --- | --- | --- |
| Anti-HLA-DR | G46-6 | PercpC5.5 | BD | 560652 | 1:80 |
| Anti-CD14 | M5E2 | PEcy7 | Biolegend | 301813 | 1:80 |
| Anti-CD16 | 3G8 | APC-H7 | BD | 560195 | 1:25 |
| Live/dead | - | eFluor506UV | eBioscience | 65-0866 | 1:833 |
| Intravenous<br>Immunoglobulin<br>(Nanogam) | - | - | Prothya<br>Biosolutions<br>Netherlands | J06BA02 | 1:100 |
| Anti-CCR2 | 48607 | AF647 | BD | 558406 | 1:5 |
| Anti-CD163 | RM3/1 | APC | Biolegend | 326509 | 1:20 |
| Anti-CD29 | MAR4 | APC | BD | 559883 | 1:5 |
| Anti-CCR4 | 205410 | APC | Novus Bio | FAB1567A-025 | 1:10 |
| Anti-C3aR1 | hC3aRZ8 | APC | Biolegend | 345805 | 1:2.5 |
| Anti-CCR5 | 2D7/CCR5 | FITC | BD | 555992 | 1:5 |
| Anti-CD18 | 6.7 | FITC | BD | 555923 | 1:25 |
| Anti-CX3CR1 | 2A9-1 | FITC | Biolegend | 341606 | 1:12.5 |
| Anti-CD11a | 345913 | FITC | Rnd systems | FAB3595F | 1:5 |
| Anti-C5aR1 | S5/1 | FITC | Biolegend | 344305 | 1:40 |
| Anti-CXCR4 | 12G5 | PE | BD | 561733 | 1:5 |
| Anti-CCR1 | 53504 | PE | Novus Bio | FAB145P-025 | 1:10 |
| Anti-CD49d | 9F10 | PE | BD | 555503 | 1:10 |
| Anti-CD11b | ICRF44 | PE | BD | 555388 | 1:10 |
| Anti-PECAM-1 | WM59 | PE | Biolegend | 303105 | 1:500 |

**Supp. Table S2:**
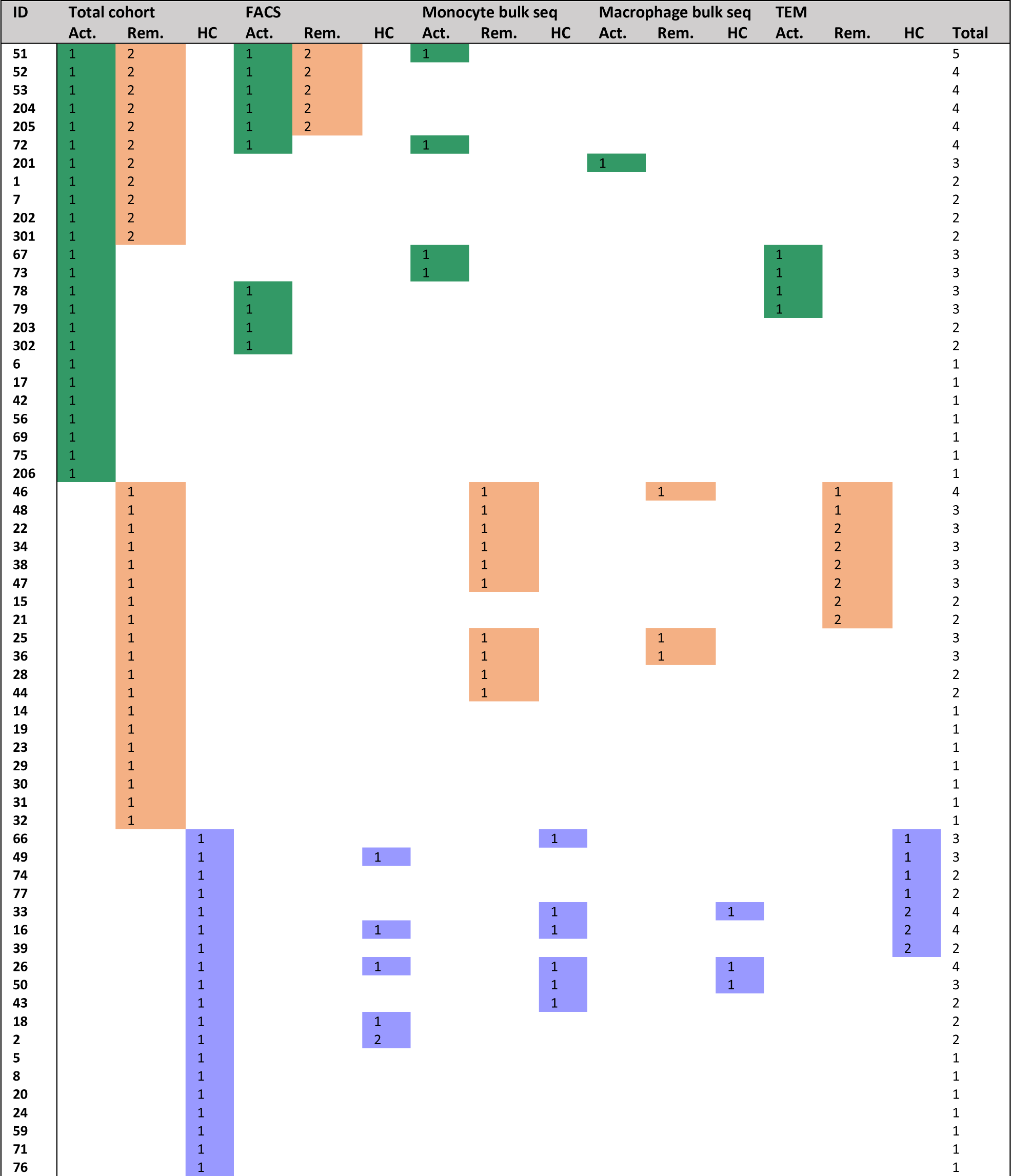
overview included patients per experiment. Act., active; Rem., remission; HC, healthy control; seq, sequencing; TEM, transendothelial migration assay

| ID | Total cohort |  |  | FACS |  |  | Monocyte bulk seq |  |  | Macrophage bulk seq |  |  | TEM |  |  | Total |
| --- | --- | --- | --- | --- | --- | --- | --- | --- | --- | --- | --- | --- | --- | --- | --- | --- |
|  | Act. | Rem. | HC | Act. | Rem. | HC | Act. | Rem. | HC | Act. | Rem. | HC | Act. | Rem. | HC |  |
| 51 | 1 | 2 |  | 1 | 2 |  | 1 |  |  |  |  |  |  |  |  | 5 |
| 52 | 1 | 2 |  | 1 | 2 |  |  |  |  |  |  |  |  |  |  | 4 |
| 53 | 1 | 2 |  | 1 | 2 |  |  |  |  |  |  |  |  |  |  | 4 |
| 204 | 1 | 2 |  | 1 | 2 |  |  |  |  |  |  |  |  |  |  | 4 |
| 205 | 1 | 2 |  | 1 | 2 |  |  |  |  |  |  |  |  |  |  | 4 |
| 72 | 1 | 2 |  | 1 |  |  | 1 |  |  |  |  |  |  |  |  | 4 |
| 201 | 1 | 2 |  |  |  |  |  |  |  | 1 |  |  |  |  |  | 3 |
| 1 | 1 | 2 |  |  |  |  |  |  |  |  |  |  |  |  |  | 2 |
| 7 | 1 | 2 |  |  |  |  |  |  |  |  |  |  |  |  |  | 2 |
| 202 | 1 | 2 |  |  |  |  |  |  |  |  |  |  |  |  |  | 2 |
| 301 | 1 | 2 |  |  |  |  |  |  |  |  |  |  |  |  |  | 2 |
| 67 | 1 |  |  |  |  |  | 1 |  |  |  |  |  | 1 |  |  | 3 |
| 73 | 1 |  |  |  |  |  | 1 |  |  |  |  |  | 1 |  |  | 3 |
| 78 | 1 |  |  | 1 |  |  |  |  |  |  |  |  | 1 |  |  | 3 |
| 79 | 1 |  |  | 1 |  |  |  |  |  |  |  |  | 1 |  |  | 3 |
| 203 | 1 |  |  | 1 |  |  |  |  |  |  |  |  |  |  |  | 2 |
| 302 | 1 |  |  | 1 |  |  |  |  |  |  |  |  |  |  |  | 2 |
| 6 | 1 |  |  |  |  |  |  |  |  |  |  |  |  |  |  | 1 |
| 17 | 1 |  |  |  |  |  |  |  |  |  |  |  |  |  |  | 1 |
| 42 | 1 |  |  |  |  |  |  |  |  |  |  |  |  |  |  | 1 |
| 56 | 1 |  |  |  |  |  |  |  |  |  |  |  |  |  |  | 1 |
| 69 | 1 |  |  |  |  |  |  |  |  |  |  |  |  |  |  | 1 |
| 75 | 1 |  |  |  |  |  |  |  |  |  |  |  |  |  |  | 1 |
| 206 | 1 |  |  |  |  |  |  |  |  |  |  |  |  |  |  | 1 |
| 46 |  | 1 |  |  |  |  |  | 1 |  |  | 1 |  |  | 1 |  | 4 |
| 48 |  | 1 |  |  |  |  |  | 1 |  |  |  |  |  | 1 |  | 3 |
| 22 |  | 1 |  |  |  |  |  | 1 |  |  |  |  |  | 2 |  | 3 |
| 34 |  | 1 |  |  |  |  |  | 1 |  |  |  |  |  | 2 |  | 3 |
| 38 |  | 1 |  |  |  |  |  | 1 |  |  |  |  |  | 2 |  | 3 |
| 47 |  | 1 |  |  |  |  |  | 1 |  |  |  |  |  | 2 |  | 3 |
| 15 |  | 1 |  |  |  |  |  |  |  |  |  |  |  | 2 |  | 2 |
| 21 |  | 1 |  |  |  |  |  |  |  |  |  |  |  | 2 |  | 2 |
| 25 |  | 1 |  |  |  |  |  | 1 |  |  | 1 |  |  |  |  | 3 |
| 36 |  | 1 |  |  |  |  |  | 1 |  |  | 1 |  |  |  |  | 3 |
| 28 |  | 1 |  |  |  |  |  | 1 |  |  |  |  |  |  |  | 2 |
| 44 |  | 1 |  |  |  |  |  | 1 |  |  |  |  |  |  |  | 2 |
| 14 |  | 1 |  |  |  |  |  |  |  |  |  |  |  |  |  | 1 |
| 19 |  | 1 |  |  |  |  |  |  |  |  |  |  |  |  |  | 1 |
| 23 |  | 1 |  |  |  |  |  |  |  |  |  |  |  |  |  | 1 |
| 29 |  | 1 |  |  |  |  |  |  |  |  |  |  |  |  |  | 1 |
| 30 |  | 1 |  |  |  |  |  |  |  |  |  |  |  |  |  | 1 |
| 31 |  | 1 |  |  |  |  |  |  |  |  |  |  |  |  |  | 1 |
| 32 |  | 1 |  |  |  |  |  |  |  |  |  |  |  |  |  | 1 |
| 66 |  |  | 1 |  |  |  |  |  | 1 |  |  |  |  |  | 1 | 3 |
| 49 |  |  | 1 |  |  | 1 |  |  |  |  |  |  |  |  | 1 | 3 |
| 74 |  |  | 1 |  |  |  |  |  |  |  |  |  |  |  | 1 | 2 |
| 77 |  |  | 1 |  |  |  |  |  |  |  |  |  |  |  | 1 | 2 |
| 33 |  |  | 1 |  |  |  |  |  | 1 |  |  | 1 |  |  | 2 | 4 |
| 16 |  |  | 1 |  |  | 1 |  |  | 1 |  |  |  |  |  | 2 | 4 |
| 39 |  |  | 1 |  |  |  |  |  |  |  |  |  |  |  | 2 | 2 |
| 26 |  |  | 1 |  |  | 1 |  |  | 1 |  |  | 1 |  |  |  | 4 |
| 50 |  |  | 1 |  |  |  |  |  | 1 |  |  | 1 |  |  |  | 3 |
| 43 |  |  | 1 |  |  |  |  |  | 1 |  |  |  |  |  |  | 2 |
| 18 |  |  | 1 |  |  | 1 |  |  |  |  |  |  |  |  |  | 2 |
| 2 |  |  | 1 |  |  | 2 |  |  |  |  |  |  |  |  |  | 2 |
| 5 |  |  | 1 |  |  |  |  |  |  |  |  |  |  |  |  | 1 |
| 8 |  |  | 1 |  |  |  |  |  |  |  |  |  |  |  |  | 1 |
| 20 |  |  | 1 |  |  |  |  |  |  |  |  |  |  |  |  | 1 |
| 24 |  |  | 1 |  |  |  |  |  |  |  |  |  |  |  |  | 1 |
| 59 |  |  | 1 |  |  |  |  |  |  |  |  |  |  |  |  | 1 |
| 71 |  |  | 1 |  |  |  |  |  |  |  |  |  |  |  |  | 1 |
| 76 |  |  | 1 |  |  |  |  |  |  |  |  |  |  |  |  | 1 |

**Supp. Table S3:** Baseline characteristics patients CD14-selected monocyte bulk mRNA sequencing. SD = standard deviation; IQR = interquartile range; BVAS = Birmingham Vasculitis Activity Score version 3; VDI = Vasculitis Damage Index; CRP = C-reactive protein; BMI = Body Mass Index; eGFR = estimated glomerular filtration rate;

|  | ANCA patients (n=14) |  | Healthy controls (n=6) |
| --- | --- | --- | --- |
|  | Active (n=4) | Remission (n=10) |  |
| **Demographics** |  |  |  |
| Age, years, mean (SD) | 57 (15) | 65 (12) | 59 (18) |
| Female sex, n (%) | 2 (50) | 2 (20) | 4 (67) |
| BMI, mean (SD) | 26 (3) | 27 (3) | 27 (7) |
| **Disease characteristics** |  |  |  |
| **Clinical subtype** |  |  |  |
| \| GPA/MPA/Renal limited, n | 0/3/1 | 8/1/1 | - |
| **ANCA subtype** |  |  |  |
| \| PR3/MPO, n | 0/4 | 5/5 | - |
| **Organ involvement** |  |  |  |
| \| Renal, n (%) | 4 (100) | 10 (100) | - |
| \| Pulmonary, n (%) | 1 (25) | 5 (50) | - |
| \| Cardiovascular, n (%) | 0 (0) | 2 (20) | - |
| \| ENT, n (%) | 0 (0) | 3 (30) | - |
| **Disease duration in days, median (IQR)** | 2 (3) | 3726 (3595) | - |
| **eGFR, (CKD-EPI), per ml/min/1.73 m<sup>2</sup>, median (IQR)** | 5 (9) | 47 (31) | 91 (40) |
| **BVAS, median (IQR)** | 13.5 (3) | 0 (0) | - |
| **VDI, median (IQR)** | 0.0 (0) | 2.5 (5) | - |
| **CRP, mg/l, median (IQR)** | 9.3 | 1.8 (4.1) | 1.5 (8.0) |
| **Monocytes, 10<sup>9</sup>/L, median (IQR)** | 0.72 (0.9) | 0.53 (0.4) | 0.51 (0.1) |
| **Use of immunosuppressive medication, n (%)** | 4 (100) | 7 (70) | 0 (0) |
| \| Glucocorticoid use, n (%) | 4 (100) | 6 (60) | 0 (0) |
| \| Prednisone dosage users, mg, median (IQR) | 1250 (0) | 5 (2) | - |
| \| Methylprednisolon <30 days, n (%) | 4 (100) | 0 (0) | 0 (0) |
| \| Azathioprine, n (%) | 0 (0) | 3 (30) | 0 (0) |
| \| Cyclophosphamide, n (%) | 1 (25) | 0 (0) | 0 (0) |
| \| Rituximab < 1 year, n (%) | 1 (25) | 3 (30) | 0 (0) |
| **Cardiovascular risk** |  |  |  |
| \| History of cardiovascular event, n (%) | 0 (0) | 4 (40) | 0 (0) |
| \| Diabetes mellitus, n (%) | 0 (0) | 2 (20) | 2 (33) |
| \| Currently smoking, n (%) | 1 (25) | 1 (10) | 0 (0) |
| \| Use of statins, n (%) | 0 (0) | 5 (50) | NA |
| \| Use of antihypertensive medication, n (%) | 2 (50) | 8 (80) | NA |

**Supp. Table S4:** Baseline characteristics patients monocyte-derived macrophage bulk mRNA sequencing. SD = standard deviation; IQR = interquartile range; BVAS = Birmingham Vasculitis Activity Score version 3; VDI = Vasculitis Damage Index; CRP = C-reactive protein; BMI = Body Mass Index; eGFR = estimated glomerular filtration rate;

|  | ANCA patients (n=4) |  | Healthy controls (n=3) |
| --- | --- | --- | --- |
|  | Active (n=1) | Remission (n=3) |  |
| **Demographics** |  |  |  |
| Age, years, mean (SD) | 54 | 55 (15) | 64 (10) |
| Female sex, n (%) | 0 (0) | 2 (67) | 2 (67) |
| BMI, mean (SD) | 23 | 25 (3) | 26 (5) |
| **Disease characteristics** |  |  |  |
| **Clinical subtype** |  |  |  |
| GPA/MPA/Renal limited, n | 1/0/0 | 3/0/0 | - |
| **ANCA subtype** |  |  |  |
| PR3/MPO, n | 1/0 | 3/0 | - |
| **Organ involvement** |  |  |  |
| Renal, n (%) | 1 (100) | 3 (100) | - |
| Pulmonary, n (%) | 1 (100) | 1 (34) | - |
| Cardiovascular, n (%) | 0 (0) | 1 (34) | - |
| ENT, n (%) | 1 (100) | 1 (34) | - |
| **Disease duration in days, median (IQR)** | 0 | 5478 (3155) | - |
| **eGFR, (CKD-EPI), per ml/min/1,73 m2, median (IQR)** | 106 | 41 (10) | 91 (8) |
| **BVAS, median (IQR)** | 27 | 0 (0) | - |
| **VDI, median (IQR)** | 0 | 5 (3) | - |
| **CRP, mg/l, median (IQR)** | 199.0 | 5.0 (0.2) | 1.5 (5.3) |
| **Monocytes, 10<sup>9</sup>/L, 10<sup>9</sup>/L median (IQR)** | 0.6 | 0.61 (0.23) | 0.61 (0.11) |
| **ANCA titer, median (IQR)** | 101 | 1.2 (1.9) | - |
| **Use of immunosuppressive medication, n (%)** | 0 (0) | 1 (34) | 0 (0) |
| Glucocorticoid use, n (%) | 0 (0) | 0 (0) | 0 (0) |
| Methylprednisolon <30 days, n (%) | 0 (0) | 0 (0) | 0 (0) |
| Azathioprine, n (%) | 0 (0) | 0 (0) | 0 (0) |
| Cyclophosphamide, n (%) | 0 (0) | 0 (0) | 0 (0) |
| Rituximab < 1 year, n (%) | 0 (0) | 1 (34) | 0 (0) |
| **Cardiovascular risk** |  |  |  |
| History of Cardiovascular event, n (%) | 0 (0) | 2 (66) | 0 (0) |
| Diabetes mellitus, n (%) | 0 (0) | 2 (67) | 1 (33) |
| Currently smoking, n (%) | 0 (0) | 1 (33) | 0 (0) |
| Use of statins, n (%) | 0 (0) | 2 (67) | NA |
| Use of antihypertensive medication, n (%) | 0 (0) | 3 (100) | NA |

**Suppl. Table S5:** Baseline characteristics patients transendothelial migration assay. SD = standard deviation; IQR = interquartile range; BVAS = Birmingham Vasculitis Activity Score version 3; VDI = Vasculitis Damage Index; CRP = C-reactive protein; BMI = Body Mass Index; eGFR = estimated glomerular filtration rate;

|  | ANCA patients (n=12) |  | Healthy controls (n=7) |
| --- | --- | --- | --- |
|  | Active (n=4) | Remission (n=8) |  |
| **Demographics** |  |  |  |
| Age, years, mean (SD) | 60 (15) | 61 (11) | 49 (21) |
| Female sex, n (%) | 2 (50) | 1 (13) | 6 (86) |
| BMI, mean (SD) | 24 (4) | 27 (3) | 24 (7) |
| **Disease characteristics** |  |  |  |
| **Clinical subtype** |  |  |  |
| \| GPA/MPA/Renal limited, n | 2/1/1 | 6/1/1 | - |
| **ANCA subtype** |  |  |  |
| \| PR3/MPO, n | 1/3 | 4/4 | - |
| **Organ involvement** |  |  |  |
| \| Renal, n (%) | 4 (100) | 8 (100) | - |
| \| Pulmonary, n (%) | 2 (50) | 2 (25) | - |
| \| Cardiovascular, n (%) | 0 (0) | 1 (12.5) | - |
| \| ENT, n (%) | 1 (25) | 4 (50) | - |
| **Disease duration in days, median (IQR)** | 0.5 (1865) | 2502 (3370) | - |
| **eGFR, (CKD-EPI), per ml/min/1.73 m<sup>2</sup>, median (IQR)** | 21 (52) | 58 (31) | 76 (30) |
| **BVAS, median (IQR)** | 12.5 (8) | 0 (0) | - |
| **VDI, median (IQR)** | 1 (4) | 2.5 (3) | - |
| **CRP, mg/l, median (IQR)** | 7.7 (18.3) | 1.6 (5.0) | 1 (6.6) |
| **Monocytes, 10<sup>9</sup>/L, median (IQR)** | 0.72 (0.8) | 0.49 (0.3) | 0.43 (0.2) |
| **ANCA titer, median (IQR)** | 80 (487) | 24 (56) | - |
| **Use of immunosuppressive medication, n (%)** | 3 (75) | 6 (75) | 0 (0) |
| \| Glucocorticoid use, n (%) | 2 (50) | 6 (75) | 0 (0) |
| \| Prednisone dosage users, mg, median (IQR) | 1250 (0) | 5 (0) | - |
| \| Methylprednisolon <30 days, n (%) | 2 (50) | 0 (0) | 0 (0) |
| \| Azathioprine, n (%) | 0 (0) | 3 (38) | 0 (0) |
| \| Rituximab < 1 year, n (%) | 1 (25) | 3 (38) | 0 (0) |
| **Cardiovascular risk** |  |  |  |
| \| History of Cardiovascular event, n (%) | 0 (0) | 1 (13) | 0 (0) |
| \| Diabetes mellitus, n (%) | 0 (0) | 2 (25) | 1 (14) |
| \| Currently smoking, n (%) | 3 (75) | 5 (80) | 4 (57) |
| \| Use of statins, n (%) | 0 (0) | 3 (38) | NA |
| \| Use of antihypertensive medication, n (%) | 2 (50) | 6 (75) | NA |

**Suppl. Table S6:**
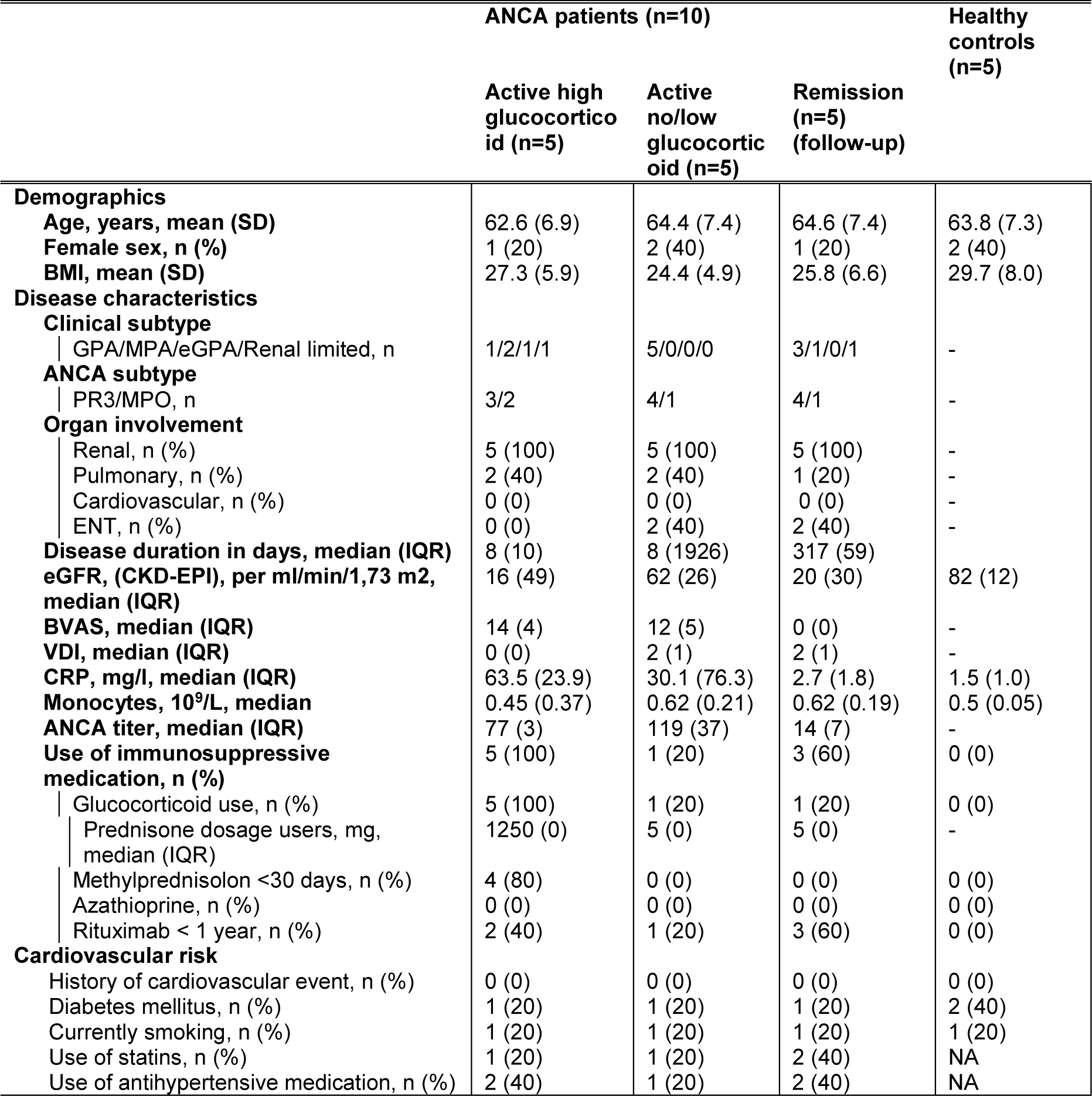
Baseline characteristics patients FACS. SD = standard deviation; IQR = interquartile range; BVAS = Birmingham Vasculitis Activity Score version 3; VDI = Vasculitis Damage Index; CRP = C-reactive protein; BMI = Body Mass Index; eGFR = estimated glomerular filtration rate;

## Supplementary data S1 – Supplementary methods

### Study subjects, samples and clinical information

To study monocyte activation and intrinsic migration capacity in AAV, blood was collected from patients with AAV (n=43) and healthy controls (n=19). Patients were either included during active disease, meaning recent AAV diagnosis or during a disease relapse, or in remission, defined by the absence of new or worse signs of disease activity and no more than 1 item indicating persistent disease activity based on the Birmingham Vasculitis Activity Score version 3 (BVASv3) (17). Patients with a clinical significant infection requiring treatment were excluded. Partners of AAV patients were invited to participate as healthy controls. Blood samples were collected either once or twice, in accordance with the timing of the patient’s hospital visits, during periods of active disease or remission. Patients were recruited from the Amsterdam University Medical Center (AUMC), Northwest Clinics and Spaarne Gasthuis Hospital in the Netherlands. On the day of blood withdrawal, information regarding demographic details, disease characteristics, and medication use was collected and a standard laboratory panel was used to evaluate kidney function, a complete blood count and the presence of inflammation. Also, disease activity and treatment or vasculitis-related organ damage was measured by the BVASv3 and Vasculitis Damage Index (VDI) (18), respectively. The blood lipid profile, including total cholesterol, LDL, HDL, and triglycerides, and use of statin and antihypertensive therapy was retrospectively collected.

### PBMC isolation peripheral blood

Blood was collected in sodium heparin tubes (Vacuette, Greiner). Using standard density gradient centrifugation (Lymphoprep, Abbott Diagnostics Technologies AS), peripheral blood mononuclear cells (PBMCs) were isolated from the blood and directly used for monocyte isolation or stored in liquid nitrogen.

### Monocyte isolation by CD14^+^ bead selection

To unravel the potential mechanisms driving differences in monocyte migration, bulk RNA sequencing was performed of CD14^+^-bead selected monocytes from patients with active (n=4) and stable disease (n=10) and healthy controls (n=6). Using freshly isolated PBMCs, CD14^+^ cells were positively selected by use of column-based immunomagnetic cell separation according to the manufacturer’s protocol (MojoSort™ Human CD14 Nanobeads, Biolegend). After monocyte isolation, the viable cell concentration was determined using a CASY cell counter (Bioké) and resuspended in RPMI1640 at a target concentration of 1 * 10^6^ viable cells/mL for further use.

### Differentiation and stimulation of monocyte-derived macrophages

To study monocyte-derived macrophage activation, we performed bulk RNA sequencing of monocyte-derived macrophages (MDMs) from patients with active (n=1) and stable disease (n=3) and healthy controls (n=3). Upon culturing, 25 million PBMCs were gently thawed in IMDM with 20% FCS and 0.1 % penicillin/streptomycin for 20 minutes. Cells were washed and resuspended in RPMI with glutamine, 10% FCS and 0.1% penicillin/streptomycin (macrophage medium) and seeded on a six well plate (6.25 million PBMCs/well). After an hour of adherence time in a 37 °C incubator, cells were washed three to four times with room temperature PBS to remove non-adherent cells. The monocytes were incubated for five days in RPMI with glutamine, 10% FCS, 0.1% penicillin/streptomycin and macrophage colony-stimulating factor (MCSF, 50 ng/ml) for monocyte-to-macrophage differentiation. On day five, cells were washed two to three times with PBS to remove non-adherent cells. MDMs were either stimulated with 10 ng/µl LPS (Escherichia coli O55:B5, Sigma) and 50 ng/µl IFNy (Biolegend), 50 ng/µl IL4 (Biolegend), 50 ng/µl IL-10 (Biolegend), or left untreated for 24 hours.

### RNA extraction

Freshly isolated CD14^+^-bead selected monocytes or *in vitro* cultured MDMs were washed twice with PBS to remove media. RNA was then isolated using RNeasy Mini Kit (Qiagen) according to the manufacturer’s instructions. An on-column DNAse treatment (Qiagen) was performed to remove genomic DNA. RNA was eluted in RNase-free H2O and yield was measured using the NanoDrop 2000 spectrophotometer (ThermoFisher Scientific). Next, the quality of the RNA samples was checked with use of the automated electrophoresis tool D1000 ScreenTape (Agilent Technologies) to ensure a RNA integrity number (RIN) score of ≥ 8.0. mRNA capture and library preparation was performed using KAPA mRNA Hyperprep (Roche). cDNA was sequenced using paired-end sequencing, with a maximum read length of 150 bp on the Novaseq 6000 (reagent kit S4.300, Illumina) at a targeted sequencing depth of 40M reads/sample.

### Data processing and analysis

Excess adapter sequences, if present, were trimmed using Trimmomatic v0.39 (parameters: “ILLUMINACLIP:./adapters_list_v6.fa:2:40:15MINLEN: 15”). Reads were aligned using HISAT2 v2.2.1 (default parameters: -p 4 –dtacufflinks–rna-strandness RF; genome: GRCh38). Counts were obtained using the Python framework HTSeq v1.99.2, and quality control was performed using FastQC v.0.11.9 and dupRadar v.1.12.1. Genes with more than 2 reads in one or more samples were kept. Count data were transformed to log2-counts per million (logCPM), normalized by applying the trimmed mean of M-values method (53), and precision weighted using voom (19). Differential expression was assessed using an empirical Bayes moderated t-test within limma’s linear model framework (20) including the precision weights estimated by voom. A second batch of monocyte RNA samples were added at a later stage, which was corrected for in the design of the analysis. Pairing of the samples (by donor) was accounted for using the duplicateCorrelation function of limma.

Geneset enrichment analysis (GSEA) was performed using CAMERA (limma package) with a value of 0.01 for the inter-gene correlation, using selected geneset collections (Hallmark collection and the BioCarta, KEGG and Reactome subsets of the C2 collection) from the Molecular Signatures Database (MSigDB; v2023.1.Hs). P-values were calculated using a two-sided directional test (direction of change, ‘up’ or ‘down’) and corrected for multiple testing using the Benjamini-Hochberg FDR. An adjusted P-value <0.1 was considered significant. Results for selected genesets and comparisons were visualized using enrichment networks. For the latter, aPEAR was used to group similar genesets together and to highlight the most important themes in the geneset enrichment results. An FDR < 0.1 was used to select genesets for the comparisons Mono_Active_MPO_vs_Control, Mono_Remission_Pooled_vs_Mono_Control and Remission_PR3_Mono_vs_MPO_Mono. Default settings for the calculation of the similarity matrix between genesets (‘jaccard’), clustering (‘markov’), and naming of the clusters (‘pagerank’) were applied. Annotation was based on the significance of the gene sets in the selected comparison illustrated by the plot.

Transcription factor activity was inferred using the decoupleR R package following the vignette for TF activity inference in bulk RNA-seq data. The TF activity scores for the chosen comparison were inferred using the CollecTRI curated collection of TFs and their transcriptional targets with the Univariate Linear Model method for TFs with at least 5 targets (54).

### Flowcytometric analysis of monocytes

To validate findings on the protein-level, flowcytometric analysis of a panel related to monocyte adhesion and migration was performed. Cryopreserved PBMCs were thawed from ANCA patients with active disease (n=10), follow-up samples during stable disease (n=5) and age-and-sex matched healthy controls (n=5). Samples from active AAV patients included equal subsets with and without concurrent high-dose (defined as ≥20 mg/day) glucocorticoid therapy. For each sample, a BD FACSCanto flow cytometer was utilized to measure six panels of migration markers, each containing intravenous immunoglobulin, a fixable viability dye eFluoro506 and monocyte subset markers (CD14, CD16, HLA-DR) in addition to 3 surface antibodies related to monocyte migration (**Supp. Table 1**). All antibodies were titrated to determine the optimal concentration using serial dilutions. For each staining reaction, in total 0.25 x 10^6^ PBMCs were incubated in a fixed staining volume of 50ul for 30 min at 4°C in the dark, and washed twice before measurements. To mitigate batch effects, we incubated all samples with a pre-made antibody cocktail and conducted measurements on the same day. Gating strategy is shown in **Supp. Fig 1**. To correct for background fluorescence, the Median Fluorescence Intensity (MFI) was subtracted with the MFI of the unstained backbone panel (including HLA-DR, CD14, CD16, viability dye). A minimum cell count threshold of 200 cells was established as a criterion for inclusion in Median Fluorescence Intensity (MFI) analyses. A Wilcoxon rank-sum test was conducted to compare the expression of migration markers on peripheral blood monocyte subsets between ANCA patients and healthy controls. Principle component analysis (PCA) was used to summarize expression of markers and to highlight differences between monocyte subsets based on their marker profile. All markers containing only positive values were selected and marker values were log-transformed to stabilize variance and improve normality and plotted using the ggfortify package. Data analysis was performed using FlowJo version 10.0.7 (FlowJo, Ashland, OR, USA) and Graphs were made using R-studio (version 2022.02.3).

### Transendothelial monocyte migration assay

To assess the adhesive and migratory capacity of CD14^+^ monocytes, a transendothelial monocyte migration assay (TEM) was performed. Human Aortic Endothelial Cells (HAEC) (Lonza) were cultured in duplo or in triplo in a 12-well plate until reaching confluency and stimulated with 1 ng/mL IL-1β (Sigma-Aldrich) and 10 ng/mL TNFα (Sigma-Aldrich) overnight for 18 hours. HAECs were gently washed to remove IL-1β or TNFα and media was changed into EGM-2 (Lonza) at the day of the assay for at least 6 hours. Next, 100,000 viable CD14^+^-bead selected monocytes were loaded onto the HAECs and gently swirled to ensure an equal monocyte distribution across the wells. The monocytes were incubated for 30 minutes at 37°C and 5% CO_2_. One follow-up experiment included a longer incubation period of 60 minutes. Subsequently, adhered and migrated monocytes were fixed by 3.7% formaldehyde (Sigma-Aldrich) and incubated for 15 minutes at room temperature. Afterwards, the medium containing formaldehyde was gradually removed and washed twice with PBS to ensure removal of non-adherent cells. Cells were imaged by a Leica DM8i Live-Cell microscope (Plan-apochromat 10/0.25 Phaco 1; Leica) and 3 pictures were taken per condition. Phase-contrast microscopy was used to identify adherent cells (light appearance, located on the HAECs layer) and transmigrated cells (dark appearance). Utilizing the NIH software ImageJ2, quantification was carried out. To control for bias, the outcome assessor was blinded for patient group.

